# Single-Cell Transcriptomics Reveals Dynamics of NK Cell Expansion in a Feeder Cell-Free Culture of PBMCs - Implications for Immunotherapy

**DOI:** 10.1101/2025.09.15.676262

**Authors:** Brian Ladd, Markella Zacharouli, Per-Henrik Holmqvist, Stefanie Renken, Pontus Blomberg, Véronique Chotteau

## Abstract

Natural killer (NK) cell therapies hold great promise for cancer treatment; however, donor-to-donor heterogeneity in the *ex vivo* expansion process remains a critical bottleneck in their supply. This study aimed to identify factors influencing donor variability in a two-week long *ex vivo* NK cell expansion from PBMCs, analyzed across three donors. Single-cell transcriptomics was applied to investigate the distribution of cell types and phenotypes, as well as trajectory inference and differential gene expression. Our results identified that several factors were associated with the variability in the final NK cell fraction and expansion, and that their influence was prevalent between culture day 3 and 8. Compared to high final NK cell fraction, a culture with low final NK cell fraction exhibited an upregulation of some stress and inflammatory genes and an increase in one specific subcluster of the NK cells already on culture day 3. It showed a low score of CD56^Bright^ CD16^−^ phenotype and high score of CD56^Dim^ CD16^+^ phenotype. It had also an increased presence of cytotoxic CD8^+^ Tm cells. Among the observed subclusters of CD8^+^ Tm cells, it exhibited a higher presence of a subcluster associated with a less differentiated and less cytotoxic phenotype as well as a lower prevalence of a subcluster associated with chemokine and cytotoxic genes. Finally, it had a major expansion of one of the CD8^+^ Tm cells subclusters annotated as NK-like T cell and characterized by a high CCR5 mRNA expression while the levels of CCL3, CCL4, and CCL5 mRNA were downregulated. The present findings point towards a potential link between CCL signaling and improved NK cell expansion performance, including possible markers for further investigations, and suggest future strategies to increase the final NK cell fraction and expansion based on donor-specific markers.

## Introduction

Natural Killer (NK) cells are cytotoxic lymphocytes of the innate immune system capable of killing target cells without prior sensitization. Due to their inherent ability to detect and lyse infected and or cancerous cells, they are an interesting approach for adoptive cell therapy. When triggered, NK cells release granules containing perforin and granzyme B to lyse the target cells [1]. Target recognition is controlled by a balance of activating and repressive ligands which is shifted in stressed or transformed cells such as tumor or virus-infected cells [2, 3]. Through the receptor CD16 they are able to recognize antibody-coated cells, and secrete cytokines that stimulate other immune cells in the tumor microenvironment [4]. At the same time, NK cells express inhibitory receptors that, among others, bind human leukocyte antigen (HLA) class 1 molecules [3]. A mismatch or loss of HLA class 1 molecules on target cells removes this inhibitory signal and promotes target cell killing, a mechanism known as “missing self” recognition [5, 6]. In line with this, it was observed that NK cells show alloreactivity against leukemic cells in the setting of haploidentical hematopoietic stem cell transplantation (HSCT) [7, 8]. Consequently, many clinical trials have evaluated the use of allogeneic NK cells derived from haploidentical donors, demonstrating a good safety profile and partially promising outcomes regarding clinical responses [9–14]. However, in solid tumors, the tumor microenvironment (TME) may render NK cells suppressed and ineffective against the tumor cells [15]. *Ex vivo* expansion of NK cells from Peripheral Blood Mononuclear Cells (PBMC) may increase the dose while also re-instating NK cytotoxic activity [16], making them good candidates for autologous immune cell therapy.

Unlike allogeneic therapies, autologous cell therapies use patient-derived immune cell that are expanded and activated *ex vivo*. An autologous immune cell product allows for infusion to the patient without immunosuppressive conditioning and risk of GvHD. Indeed, a phase I study (NCT04558853) of *ex vivo* expanded autologous NK cells from PBMCs in Multiple Myeloma (MM) showed good safety and tolerability [17]. Initial clinical trials using autologous NK cells have demonstrated successful engraftment of infused cells however without mediating remarkable tumor regression [18–20]. For this reason, NK cells are envisioned in combination with other immunotherapies, such as monoclonal antibodies. To this end a phase 2 study (NCT04558931) with autologous NK cells in combination with Isatuximab, a monoclonal antibody targeting CD38 is underway. In these studies, NK cells were expanded from patient-derived PBMCs using wave bioreactor resulting in a cell product enriched in highly active NK cells[17, 21]. Alternatively, the field is using NK cell potential by developing approaches such as CAR-NK cells [4, 22].

One of the challenges with an autologous product is the limitations associated with the quality and quantity of the starting material, with both the expansion rates and the fraction of NK cells in the final product subject to variations between donors [21]. The patient health or other aspects of biological variation may influence the composition and responsiveness of PBMCs used for expansion. To improve the robustness of this expansion and reduce the heterogeneity of the final cell products, it is important to identify parameters that could lead to these donor-to-donor variations. The *ex vivo* cultivation of NK cells is a complex process requiring the correct balance of various activation and inhibitory stimuli. Those include the addition of cytokines from the interleukin family, such as IL-2, as well as the addition of feeder or accessory cells. A manufacturing benefit of an autologous NK cell therapy approach is the ability to use the donor’s own cells as a source of accessory cells which can greatly improve the expansion of NK cells compared to cytokines alone [23]. This stimulatory effect comes from both secreted factors and cell-to-cell contacts with the accessory cells. Many different cell types within the PBMC fraction, including T cells and monocytes, coordinate to increase the expansion of NK cells *ex vivo* [24, 25]. Due to their highly heterogenous nature, PBMCs are challenging to use, since there is variation between donors. Understanding and controlling this variation can provide higher consistency of a simplified manufacturing process without cell selection or feeder cells [17, 21].

Single cell transcriptomics (scRNA-seq) has revolutionized the study of cellular heterogeneity, particularly in complex systems, such as the immune system. The ability of scRNA-seq to provide gene expression at the single cell level allows for the identification of new and distinct subpopulations and indicators of phenotypes that have been previously masked in bulk RNA sequencing. Thanks to this level of detail, investigators have been able to better understand the heterogeneity within the PBMC fraction [26]. This ability to resolve fine details and to classify cells into unique subpopulations makes scRNA-seq an ideal tool to understand the complex *ex vivo* expansion process of NK cells issued from a PBMC culture. However, we have not found any previously published reports on the use of scRNA-seq to better understand the donor-to-donor heterogeneity of an *ex vivo* NK cell expansion culture process.

In the present study, we have studied by scRNA-seq donor derived PBMCs before and during *ex vivo* NK cell expansion to identify factors responsible for donor-to-donor heterogeneity. The isolated PBMCs were cultured in the presence of interleukin 2 (IL-2) and activated by the anti-CD3 antibody, OKT3, in a scheme known to preferentially expand and reactivate NK cells [16]. In addition to NK cells, the final product also contained T cells to a variable and donor dependent degree. Three donors, one healthy and two MM patients, were chosen based on observed differences in NK cell expansion. The cells from the three expansions (donors) were analyzed at four timepoints (day 0, 3, 8, 15) during expansion by flow cytometry as well as single cell sequencing.

## Materials and Methods

### PBMC culture and NK cell expansion

The PBMCs used in this study were provided by XNK Therapeutics, and cultured according to a previously described expansion protocol [21]. In brief, the PBMCs were isolated by density gradient centrifugation using Ficoll (Cytiva), washed and frozen in aliquots. Thawed cells were cultured in Cellenix SCGM serum-free medium (CellGenix, Freiburg, Germany, now owned by Sartorius Göttingen, Germany) supplemented with 5% human serum (Access Biologicals, Vista, CA, USA), 500 IU/mL recombinant human inerlukin-2 (IL-2) (Proleukin, Novartis Pharmaceuticles, East Hanover, Nj, USA). Monoclonal anti-CD3 antibody (Orthoclone OKT-3) (Ortho Biotech, Raritan, NJ, USA) was added at the beginning of the culture to a final concentration of 10 ng/mL, and was not added during subsequent medium additions. For each donor, the PBMCs were seeded at a concentration of 0.5 × 10^6^ cells/mL in multiple T-25 flasks (Nunc), and expanded by splitting and re-seeding them at this concentration every 2- or 3-days. On days 3, 8, and 15 one of the flasks per donor was harvested and cryopreserved for further analyses.

### Flow cytometry

The cells were analyzed by flow cytometry on days 0, 3, 8, 10, and 15 using standard procedures with fluorochrome conjugated monoclonal antibodies against CD56 and against CD3. The cells were first washed with PBS, and then stained with live/dead fixable Aqua (Invitrogen, Carlsbad, CA, USA), CD3-PE (BD Biosciences ref 345765), and CD56-APC (BD Biosciences ref 341027), at 4° C for 30 min. For the day 0 sample, CD14, CD19 and CD45 were also included to analyze the more complex composition of PBMCs. The labeled cells were washed once with PBS the fixated in 4% BD CellFIX (BD Biosciences). Data acquisition and analysis were done with a FACSVerse flow cytometer (BD Biosciences, San Jose, CA, USA) and FlowJo software (Treestar Inc., Ashland, OR, USA). A Side Scatter (SSC)/Forward Scatter (FSC) gate was used to distinguish the cells from debris, and FSC-A vs FSC-H to select single cell events. LIVE/DEAD Fixable Aqua negative cells were gated as live cells. NK cells were gated as the CD3^−^ CD56^+^ population. NK-like T cells and T cells were gated as CD3^+^ CD56^+^ and CD3^+^ CD56^−^ populations, respectively. All three populations were given as fractions of live cells.

### Single-cell RNA capture, library construction, and sequencing

For each donor, the four samples, corresponding to the four time points (day 0, 3, 8, and 15), were thawed in a water bath at 37° C before multiplexing with the Single-cell multiplexing kit from BD according to the manufacturer’s instructions (Doc ID: 210970) (BD Bioscience). Due to the presence of red blood cells in the day 0 samples, a red blood cell lysis was performed with RBC lysis buffer (Cat Num: 00-4333-57) according to the protocol provided by BD (Doc ID: 210964). The isolation of single cells, subsequent capture of the RNA, and then cDNA synthesis was performed with the BD Rhapsody Express (BD Biosciences) according to the manufacturer’s protocol (Doc ID: 210967) using one cartridge per donor. A target of 5000 cells per time point was used, which resulted in a total of 20000 cells per cartridge.

The Human Immune Response Panel (Cat Num: 633750), which amplifies a set of 399 genes relating to immune cells, was used to prepare the libraries from the cDNA containing BD Rhapsody beads (Doc ID: 23-24122). The libraires were each given a unique primer to be later identified. The library quality and fragment length were checked with capillary gel electrophoresis on a Bioanalyzer 2100 (Agilent Technologies, Santa Clara, California). The high sensitivity DNA analysis kit was used according to the manufacturer’s protocol (Cat Num: 5067-4626, Agilent Technologies, Santa Clara, California).

After library preparation the three libraries were pooled together and sequenced on a Novoseq P150 with 200M reads per library for the RNA samples and 13.33M reads for each sample multiplexing library.

### Pre-processing

The alignment of the sequencing data was done on the cloud-based bioinformatic platform SevenBridges (Charlestown, MA, USA). The BD Rhapsody™ Targeted Analysis Pipeline Version 1.11 was used to align the FASTQ files from sequencing, generate the count matrices, and provide an initial cell type prediction.

Quality control and pre-processing were performed using the Seurat library in R (Hao et al. (2021), version 4.2.2). Low-quality cells were removed based on an RNA feature and total RNA count thresholds that have been previously reported, cells were removed if they had less than 60 or more than 200 unique genes (features) or if the total number of RNA counts was above 5000 [27]. Additionally, cells that are labeled as “Multiplet” or “Undetermined” by the alignment pipeline were also removed. The data were also normalized, scaled, and highly variable features were identified using the functions NormalizeData, ScaleData, and FindVariableFeatures.

### Dimensionality reduction and clustering

Principal Component Analysis (PCA) was performed on the overall data using the function RunPCA, with the features set to the variable features found in the previous step. The Seurat object was then subsetted based on cell type, followed by clustering.

### Clustering resolution

To determine the optimal clustering resolution the package ChooseR was used [28]. This package provides a framework to test different clustering parameters and outputs metrics that can be used to score and choose the best parameter values based on robustness. Additionally, this package supports a data driven approach to select parameter values that allows an objective parameter estimation step. The resolution parameter in the Seurat function “FindClusters” was varied by ChooseR and the following values were tested (0.1, 0.2, 0.3, 0.5, 0.8, 1, 1.2, 1.6). Silhouette analysis was used to score the selected resolutions and the value that had the lowest silhouette score was chosen.

### Cell cluster annotation

Cell clusters were annotated using marker genes known to be associated with particular cell types. The following datasets containing maker genes and cell labels were used: MonacoImmuneData (MID) [29], Human Primary Cell Atlas Data (HPCA) [30, 31], sc-Type [32], BlueprintEncodeData (BE) [31, 33, 34], Database Immune Cell Expression Data (DICE) [35] and azimuth with reference PBMC l2 [36]. The cell types were initially determined by the BD Rhapsody™ Targeted Analysis Pipeline Version 1.11 and were later refined based on the consensus among all cell type annotation methods.

### Trajectories

The trajectories were determined using the Monocle3 package [37–41]. The NK cells were subsetted prior to running the trajectory inference pipeline. Forty-three PC dimensions were used in the UMAP visualizations. This number was chosen based on the criterion that at least 90% of the variance in the data should be explained and that the percent change in variation between consecutive PC dimensions should be less than 5%. A resolution of 6e-4 was used in the function *cluster_cells*. The starting node was chosen based on the location of the cells from day 3.

### Cell-type fractions

The fractions of each cell type and subcluster were calculated by summing all the cells labeled with a specific cell type or subcluster, for each donor and sampling time point. This was then divided by the total number of cells for that cell type or all the cells for that donor and sampling time point. It was finally multiplied by 100 to get the percentage of each cell type. The following formula used was:

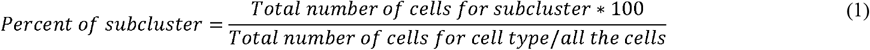

### Module scores

The module scores were calculated using the *AddModuleScore* function in Seurat with the parameters, *ctrl* and *nbin* set to 30 and 10 respectively.

### Statistical significance testing

Violin plots with statistical significance testing were generated using the function *stat_compare_means* with default parameters from the package *ggpubr*. The statistical test used was the Wilcoxon signed-rank test.

### Differential gene expression

Significantly differentially expressed genes between subclusters or donors were found by using the function *FindAllMarkers* in Seurat with default parameters. The results were filtered to include only genes that had an adjusted p value, *p_adj*, larger than 0.05.

## Results

### *Ex vivo* expansion of PBMCs revels key time points to discriminate favorable expansion outcomes

The cell types of the *ex vivo* expanded PBMCs were evaluated by flow cytometry throughout the culture as described in the Materials and Methods Section. The ending fold NK cell expansion and ending fraction of NK cells is shown in Table 1.

**Table 1:**
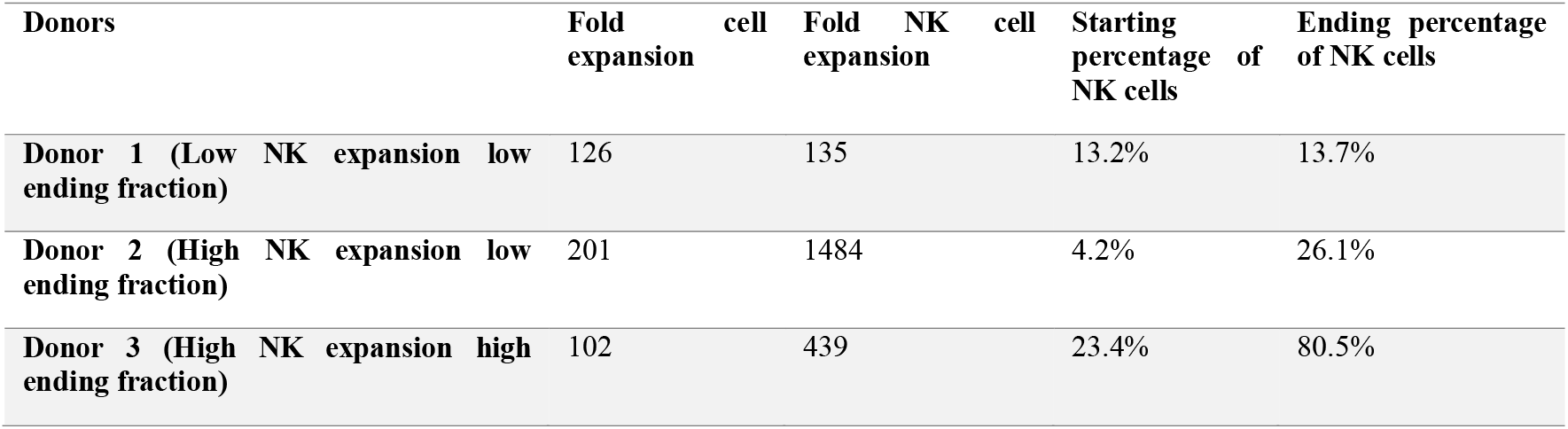
Fold expansion and ending NK cell fraction as determined by flow cytometry for the *ex vivo* cultivation of the PBMCs from the three donors.

Donors 2 and 3 were patients diagnosed with multiple myeloma while donor 1 was a healthy donor. The three donors were selected because they had very different expansion patterns that covered the range typically observed when using this *ex vivo* expansion process. Donor 1 had a low fold cell expansion (135) and a low ending fraction of NK cells (13.7%). Donor 2 had a high fold cell expansion (1484), but, however, a low ending fraction of NK cells (26.1%). Donor 3 represents the ideal case of the *ex vivo* process with a high fold cell expansion (439) and high ending fraction of NK cells (80.5%). The fold expansion of NK cells and apparent growth rate of the expansion process can be seen in Figure 1A and 1D, with the fold expansion of NK cells being determined with equation 2.

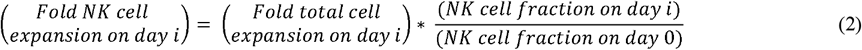

**Figure 1:**
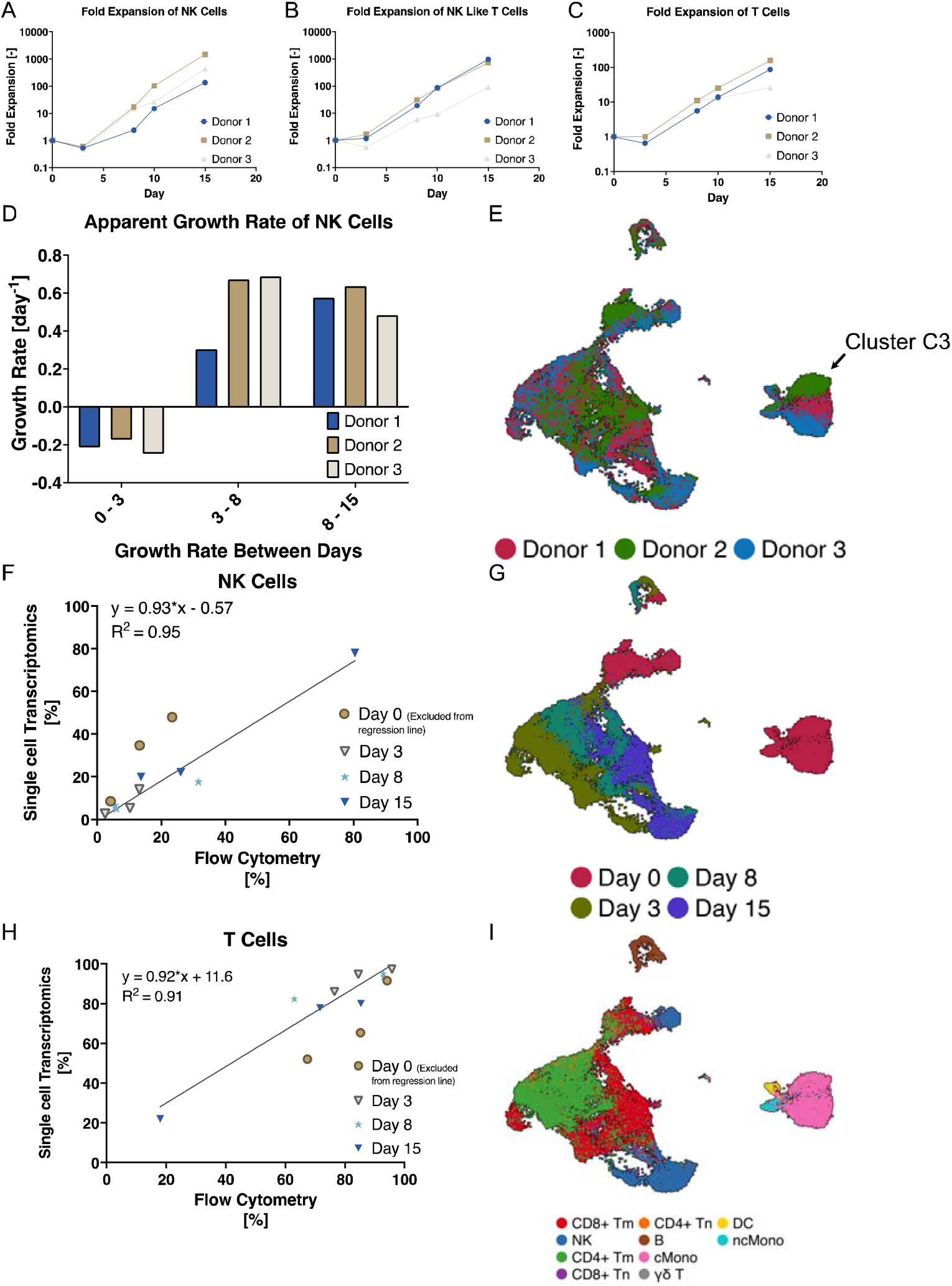
A) Fold expansion of NK cells (CD56^+^, CD3^−^) of the different donors on each of the days measured by flow cytometry. B) Fold expansion of NK like T cells (CD56^+^, CD3^+^) of the different donors on each of the days measured by flow cytometry. C) Fold expansion of T cells (CD56^−^, CD3^+^) of the different donors on each of the days measured by flow cytometry. D) Apparent growth rate measured by flow cytometry between days 0 - 3, 3 - 8, and 8 - 15. E) UMAP projection of all the cells at all the analyzed days from each donor colored by donor, where cluster C3, indicated by an arrow, is the only region where the cells are clustering based on donor. F) Comparison between cell type determination for NK cells based on flow cytometry and transcriptomic annotation. G) UMAP projection of all the cells from each donor colored by day of analysis. H) Comparison between cell type determination for T cells based on flow cytometry and transcriptomic annotation. I) UMAP projection of all the cells at all the analyzed days from each donor colored by predicted cell type.

Similarly, to Table 1, Figure 1A shows that donor 1 had the poorest NK expansion performance, measured by the fold change, while donors 2 and 3 had a better NK expansion. Figure 1A shows that, between days 3 and 8, donor 1 diverged from the other donors. Between days 8 and 15, all the donors had a similar NK cell fold change, however, donor 3 had a slightly lower fold change between days 8 and 10. As shown in Figure 1D, the slow growth rate between days 3 and 8 of donor 1 was responsible for the lower fold expansion by day 15, since the growth rates between days 8 and 15 were roughly similar among the donors.

Whereas donors 2 and 3 had a similar NK cell expansion, donors 1 and 2 had a similar T cell expansion. Figures 1B and 1C show the fold expansion of the NK-like T cells and T cells respectively. The growth behaviors of the NK-like T cells and T cells of donors 1 and 2 were very similar, while a diverging pattern for donor 3 was observed for the NK-like T cells between days 0 and 3 and after day 8 for the T cells.

Based on the cell fold expansion data determined by flow cytometry, days 0, 3, 8 and 15 stood out as important time points. Either at these time points or in between them, the *ex vivo* cultures diverged from each other in all the measured cell types. In addition to these changes these time points also corresponded to important steps in the process; i.e., day 0 corresponded to the starting material; day 3 was shortly after the T cell activation by OKT3; day 8 was in the middle of the expansion process where the NK cell growth rate was at its peak; and, lastly, day 15 was the end of the process representing the final product. Therefore, it was decided to elucidate the mechanisms of the observed divergences, i.e., donor-to-donor variations, by analyzing the cell transcriptome at these time points. Each of these samples included many cell types with inherent heterogeneity within each cell type and was thus highly heterogenous. Compared to bulk RNAseq, which would have required laborious cell sorting representing as well a supplementary source disturbance on the cells, single cell transcriptomics (scRNAseq) offered a powerful approach with no need of cell sorting. Additionally, this provided a highly sensitive analysis enabling the identification of small changes without necessitating a large number of cells.

### Limited batch effects observed in the single cell transcriptomic data sets

To minimize the batch effects of the single cell sequencing, for each donor the four selected analysis time points (days 0, 3, 8, 15) were multiplexed together using the Human Multiplex kit from BD. After multiplexing, the cells were captured on one BD Rhapsody cartridge per donor. The subsequent library preparation was done separately for each cartridge. After the library preparation, the libraries were pooled together and run on a single flow cell of an Illumina sequencer.

The raw sequencing output was aligned using a pipeline provided by BD on the cloud computing service SevenBridges. The output of the alignment was then processed in R using the Seurat package. Figure 1E shows a UMAP projection of the cells from all the analyzed days highlighted by donor and similarly Figure 1G shows at which time point each cell originated from.

As can be seen in Figure 1E, the cells did not cluster based on donor, except for a small cluster on the middle right of the figure, denoted “cluster C3” (see arrow), including separately all the 3 donor cells. When looking at Figure 1G it can be seen that cluster C3 originated entirely from day 0. It can also be seen in Figure 1G that the cells tended to be clustered based on the analysis time point.

The alignment pipeline could also give a prediction of the cell type for each cell. This was accomplished through an automatic annotation process using the marker genes listed in Table S1. Figure 1I shows the UMAP projection with the cells colored based on the predicted cell type.

From Figures 1E and 1I, it was concluded that cluster C3 donor annotated mostly to classical monocytes, with small numbers of other cell types. During the culture, it is known from previous studies and in house observations that the monocytes are usually lost by day 3, but these might have a significant initial impact in triggering lymphocyte proliferation.

In summary, Figures 1G and 1I show that the cells were primarily clustered by the sample time point and the identity of the cell type. The clustering based on donor appeared to be small, i.e., limited to cluster C3. Notice as well that since the libraires for each donor were prepared independently there was also some batch-to-batch variation when analyzed in the UMAP projection.

### Cell type comparison of flow cytometry and single cell transcriptomics

The cell types determined via annotation of the scRNAseq data were compared to the cell types determined through flow cytometry. The scRNAseq did not have a gene signature to determine NK-like T cells so cells that expressed both CD56 and CD3 were added to the T cell fraction in the flow cytometry data, as the NK-like T cell phenotype is more closely related to a T cell than an NK cell. Figure 1F shows the percentage of total cells that were NK cells for each day and donor as determined by flow cytometry and scRNAseq on the x- and y-axis respectively; and likewise Figure 1H shows T cells. The line of best fit was determined by linear regression in GraphPad Prism where the day 0 samples had been excluded. These samples were excluded from the fit determination due to the very large proportion of monocytes, and due to the fact that the transcriptomics data were very different from the other time points, with a low number of genes and mRNA transcripts per cell using the Human Immune Response Panel array of BD. Figures 1F and 1H shows a good fit between the flow cytometry and scRNAseq both for the NK cells and the T cells, indicating a good alignment of the phenotype and the transcriptome.

### Sub clustering of the cell types reveals a transcriptomic shift of NK cells during *ex vivo* expansion

The high dimensionality of scRNA-seq data enabled further subdivision of the cells beyond a single cell type, facilitating the identification of subclusters that reveal distinct phenotypes, such as NK-like T cells or the CD56 Dim and Bright NK cell subsets. The cells from all the donors were first separated based on the cell type determined during the alignment process. These cell types were then further clustered using a separate optimal resolution parameter. This parameter was chosen based on the average silhouette score from multiple runs. This allows for a data-driven selection of important parameter(s).

If the initial cell type annotation algorithm erroneously classified the cell type of some cells, the subclustering could generate some cell subclusters that were missing the parent clusters cell type. To investigate this, the subclusters from each cell type were annotated by six other cell type annotation databases, giving a total of seven annotation databases. Figure S1 shows the consensus of three cell types for the seven annotation databases. If the cell type predicted by one of the new databases was not in the original 10 cell types it was labeled as “other”.

Figure S1 shows that there was a high consensus among the different databases. The subclusters of cells that had a low degree of consensus or a different cell type than the parent cluster typically had very low cell numbers. Such consensus was observed for all the cell types except the dendric cells which mostly annotated to monocytes.

One of the databases, scType, contained an annotation signature for NK-like T cells. When using this database, the CD8^+^ Tm-0 cell subcluster was annotated to NK-like T cells.

The subclustering allows for the identification of unique populations of cells that are present at different time points during the *ex vivo* expansion process. As was seen in the UMAP projection in Figure 1G the cells tended to be clustered by analysis day. This indicated that the transcriptomic signature of the cells was uniquely different between each time point. In other words, within one cell type, there were different groups of cells that were present at each time point. Figure 2A-2F1 shows the time course abundance of the different subclusters of NK cells for each donor, calculated in relation to the total of all the cells (Figure 2A, 2B, 2C) and in relation to all the NK cells (Figures 2D, 2E, 2F), with equation 2. The area of each color represents the percentage of total cells for a particular subcluster. For example, on day 0 all the donors had exclusively the NK-1 subcluster, with a small percentage <20% of the total cells for each donor on that day. Likewise on day 15, donor 3 had close to 80% NK cells with roughly 60% of these being the NK-0 subcluster.

**Figure 2:**
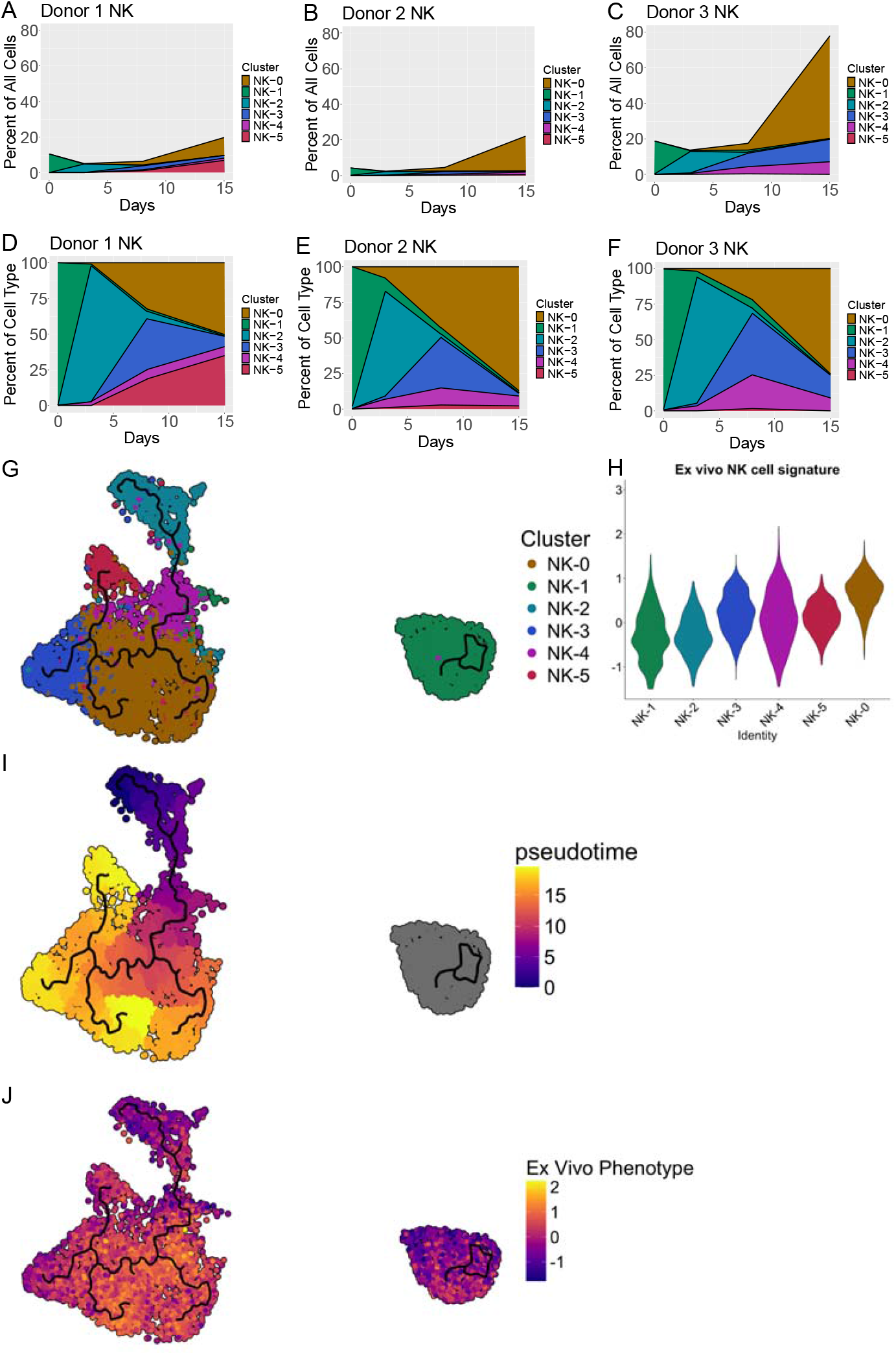
A - C) Percentage of each subcluster of NK cells with respect to the total cells on each day for each donor during the ex vivo expansion process. D - F) Percentage of each subcluster of NK cells with respect to only the NK cells on each day for each donor during the ex vivo expansion process. G) Trajectory inference projected onto a UMAP plot of all NK cells colored by NK subcluster. H) Module score of the ex vivo phenotype for the most prevalent NK cell subclusters. Subcluster NK-0 has a significantly higher module score compared to all other subclusters (comparisons not shown for clarity, p values are given in Table S3) I) Trajectory inference projected onto a UMAP plot of all the NK cells colored by pseudotime. J) Trajectory inference projected onto a UMAP plot of all the NK cells colored by *ex vivo* gene signature (the higher the score, the closer to the *ex vivo* phenotype).

For all the donors, the subclustering followed a similar pattern of NK cell expansion characterized by an initial NK cell population made up exclusively of the NK-1 subcluster. NK-2 subcluster dominated the culture on day 3, i.e., when the early activation step took place. Then on day 8, during the rapid expansion phase, the NK-2 subcluster split into mostly the NK-0 and NK-3 subclusters. By day 15, the majority of the NK cells were found in the NK-0 subcluster, with donor 3 characterized by a small amount of subcluster NK-3 remaining. For donor 1, which had a reduced NK cell expansion, a significant amount of subcluster NK-5 occurred on day 15. This temporal trend in subcluster development is depicted in greater detail with the trajectory inference in Figure 2G.

The trajectory inference, Figure 2G, reveals that subcluster NK-1 as well as the NK cells on day 0, were isolated from the other subclusters. It goes on to reveal that all the other subclusters originated from NK-2, from which they then all pass through subcluster NK-4, a dynamic that was not fully captured in Figures 2A-2F. Subclusters NK-5, NK-3, and NK-0 were all terminal subclusters indicating that once NK cells differentiated into them, they did not transition to a different subcluster.

The cells of donor 1 had a different subcluster evolution than the ones of donors 2 and 3 as can be seen in Figures 2A and 2D. The donor 1 culture had also a lower fold expansion of NK cells compared to donors 2 and 3 as observed in Figure 1A, with the key contributing factor being a slow growth rate between days 3 and 8. The growth rate after day 8 was roughly similar among all the donors (Figure 1D). This indicated that the different NK cell subclustering trends, notably the formation of subcluster NK-5, may not be associated to the slow growth rate but that this was rather due to another effect prior to day 8 in the culture. This effect can be seen when comparing the gene expressions of NK cells on day 3 in the *ex vivo* expansion process issued from the different donors. The heatmap in Figure 3A shows the expression of the significantly differentially expressed genes that were common to both donor 1 and donor 2, and to donor 1 and donor 3 on day 3. Table 2 groups the genes into classes and describes briefly their function.

**Table 2:**
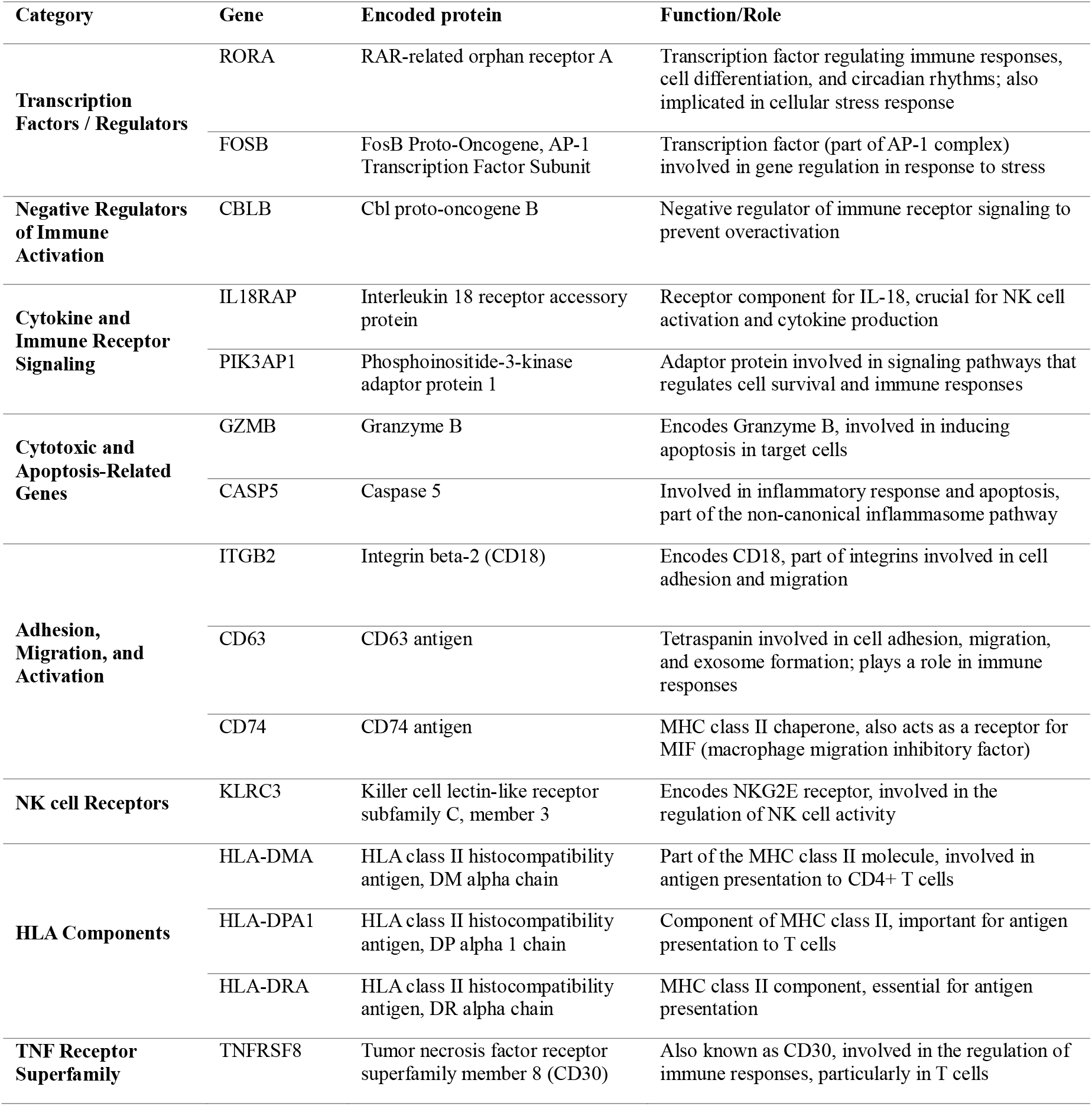
Gene descriptions of the significantly differentially expressed genes that were common to both donor 1 vs donor 2, and donor 1 vs donor 3 on day 3.

**Figure 3:**
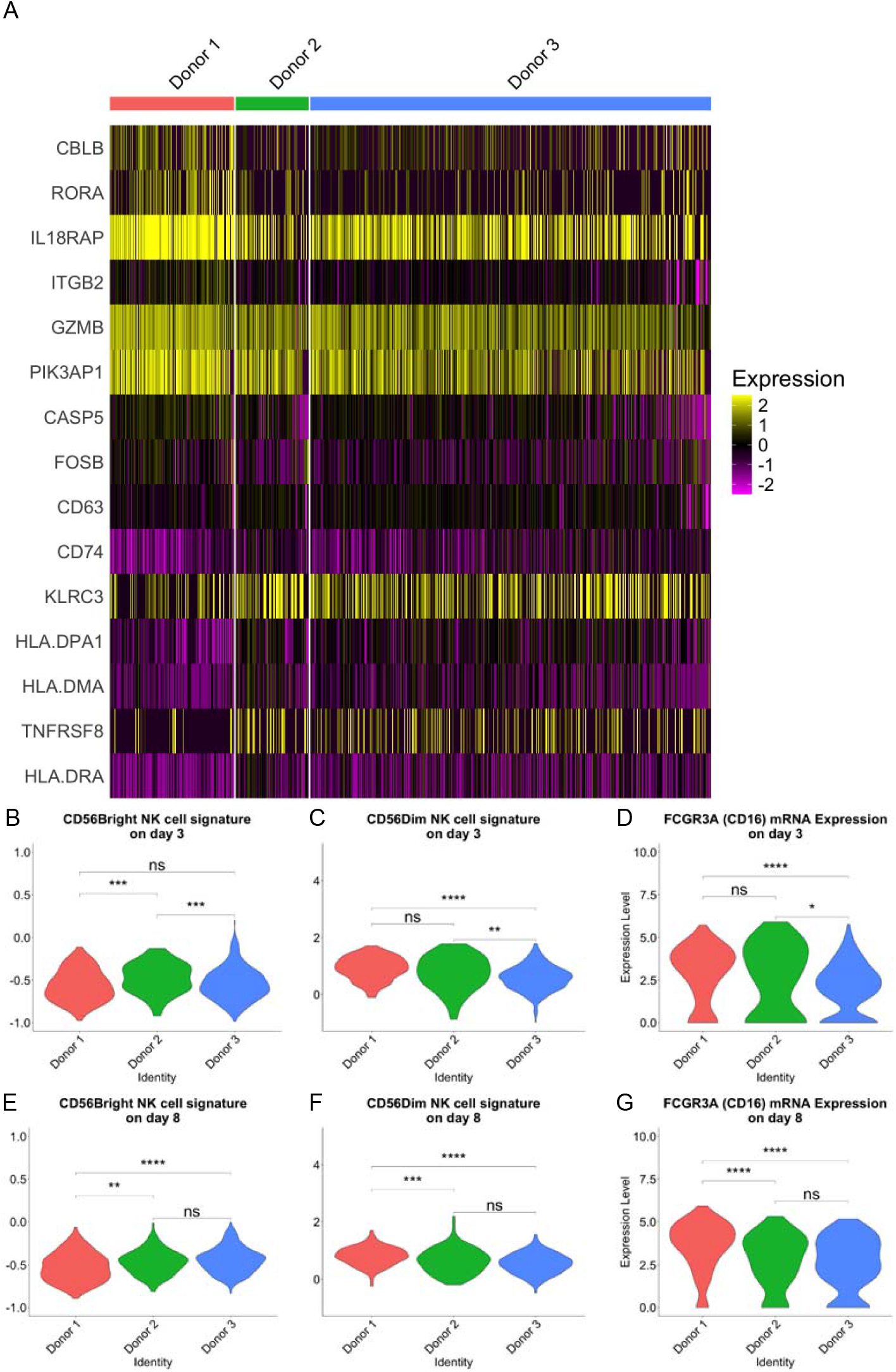
A) Heatmap of the gene expression levels of the significantly differentially expressed genes that were common to both donor 1 and donor 2, and donor 1 and donor 3 on day 3. B) CD56^Bright^ CD16^−^ NK cell signature for the NK cells of all the donors on day 3. C) CD56^Dim^ CD16^+^ NK cell signature for the NK cells of all the donors on day 3. D) Expression of CD16 mRNA in NK cells on day 3 of the ex vivo expansion. E) CD56^Bright^ CD16^−^ NK cell signature for the NK cells of all the donors on day 8. F) CD56^Dim^ CD16^+^ NK cell signature for the NK cells of all the donors on day 8. G) Expression of CD16 mRNA in NK cells on day 8 of the ex vivo expansion.

On day 3, the NK cells of donor 1, exhibiting the slowest growth rate in comparison with the other donors, showed an upregulation in the transcripts related to inflammatory environments (CASP5, IL18RAP), damage and stress (FOSB, RORA [42]), as well as in the transcripts involved in the negative regulation of NK cells such as CBLB [43].

The two major subsets of NK cells, phenotypes CD56^Dim^ CD16^+^ and CD56^Bright^ CD16^−^, differ in many aspects, such as the cytotoxic capacity. However, it has also been shown that their proliferative capacity also differs, with the CD56^Bright^ CD16^−^ phenotype having a higher growth rate [44]. The signatures (genes listed in table S1) of each of these phenotypes along with the gene expression of CD16 are shown for days 3 and 8 for all the donors in Figures 3B – 3G.

On days 3 and 8, the cells of donors 2 and 3 had a higher gene signature associated with the CD56^Bright^ CD16^−^ phenotype than donor 1. Likewise, the cells of donor 1 had a higher CD56^Dim^ CD16^+^ phenotype for days 3 and 8 than the other donors. The expression of CD16 mRNA followed the same trend where the cells of donor 1 had an upregulation of CD16 and days 3 and 8 compared to the other donors. It has been shown previously that when using this expansion protocol, a third population arise with features of both CD56^Bright^ and CD56^Dim^ cells [45].

Figure 2D – 2F shows that the NK subcluster NK-0 represents the majority of the NK cells at day 15, and that it is the subcluster responsible for the majority of the NK cell fold change in the *ex vivo* expansion process. The increase in NK-0 cell fraction is a combination of growth and NK cells from other subclusters differentiating into the NK-0 subcluster, as can be seen in Figure 2G, considering the pseudotime evolution given in Figure 2I. A measurable and distinct phenotype of NK cells, detectable by flow cytometry, has been previously described for the *ex vivo* expansion method used here [45]. This phenotype is characterized by an increase in the surface expression of several important receptors for NK cells, listed in Table S2. Since the NK-0 subcluster represents the majority of the NK cells at the end of an expansion in culture, it was interesting to evaluate how well this subcluster showed this described phenotype in comparison with the other NK subclusters. Figure 2H shows the module score using the genes of the *ex vivo* phenotype, listed in Table S2, for all the subclusters of NK cells, showing a significantly higher module score for NK-0. This higher module score showed that subcluster NK-0 had the gene expression pattern most closely resembling this phenotype compared to the other subclusters (Figure 2H). Notably subclusters NK-3, NK-4, and NK-5 had also an elevated module score compared to subclusters NK-1 and NK-2, but not to the same extent as NK-0. This is also visualized in Figure 2J, where it can be seen that the area with the strongest *ex vivo* phenotype signature is associated to NK-0 subcluster and to high pseudotime values.

### Cytotoxic CD8^+^ Tm cells influence NK cell expansion outcomes

A dominance of some subclusters evolving with time observed for the NK cells was also identified for the CD8^+^ Tm cells. Figures 4A – 4F show the results of a study of the evolution with time of the CD8^+^ Tm cells, performed similarly to the identification of NK-0 to NK-5 subclusters (Figures 2A – 2F). It can be seen that the total percentages of T cells for donors 1 and 2 were more similar to each other than to donor 3, corroborating the flow cytometry data of T cells shown in Figure 1C.

**Figure 4:**
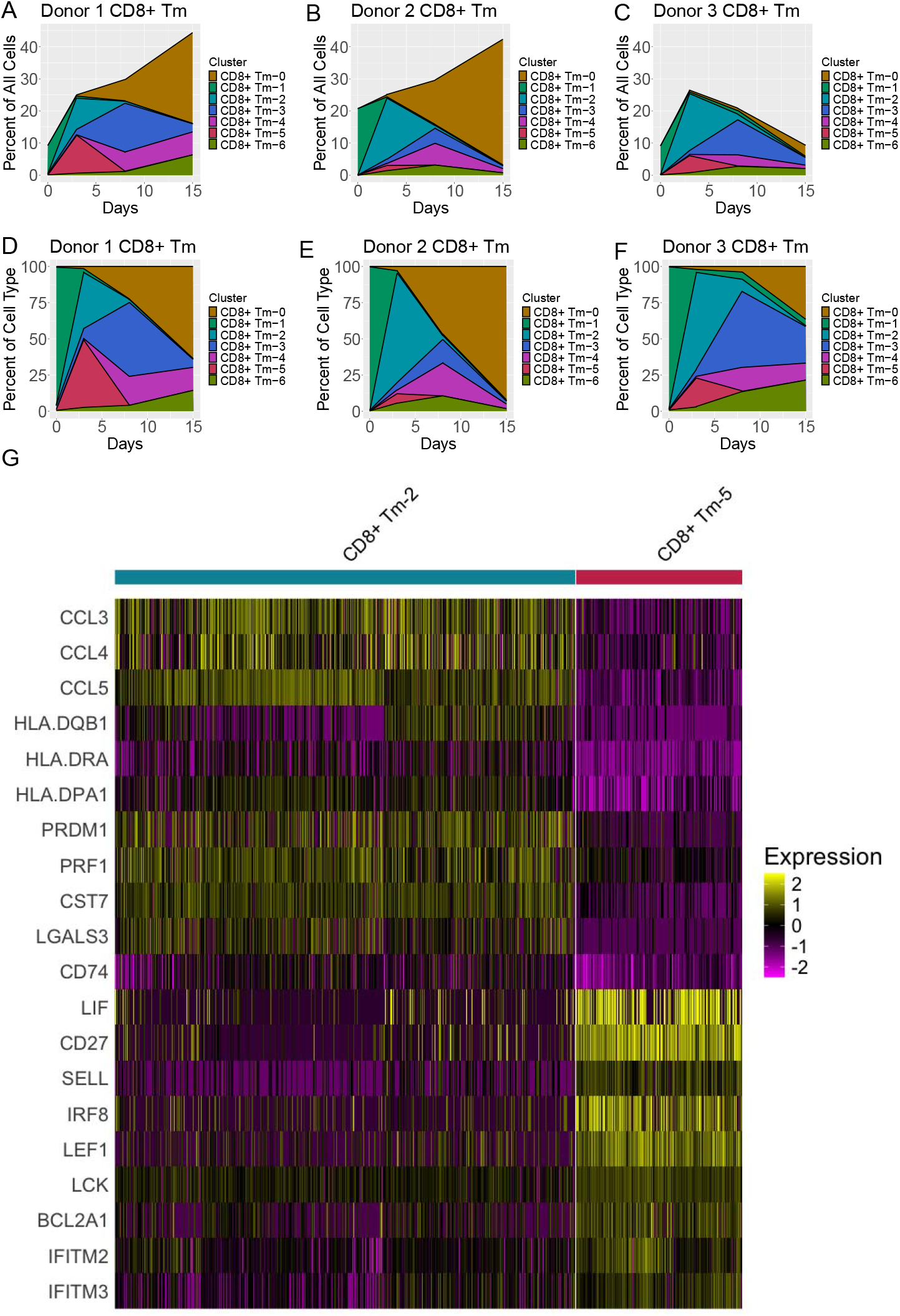
A - C) Percentage of each subcluster of CD8^+^ Tm cells with respect to the total cells on each day for each donor during the ex vivo expansion process. D - F) Percentage of each subcluster of CD8^+^ Tm cells with respect to only the CD8^+^ Tm cells on each day for each donor during the ex vivo expansion process. G) Heatmap of the gene expression of the top 10 highest fold change genes between clusters 2 and 5 of the CD8^+^ Tm cells of all the donors and all the time points.

On day 0, the CD8^+^ Tm cells fell into a single subcluster, CD8^+^ Tm-1, for all the donors. Subsequently, on day 3, the subcluster CD8^+^ Tm-2 was the most prevalent for donors 2 and 3, while, for donor 1, subcluster CD8^+^ Tm-5 was equally large as subcluster CD8^+^ Tm-2 and was larger than donors 2 and 3. By day 8, the proportion of subcluster CD8^+^ Tm-0 began to increase for donors 1 and 2 but not for donor 3, likewise subcluster CD8^+^ Tm-3 for donors 1 and 3 (but not for donor 2). Finally, subcluster CD8^+^ Tm-0 was dominating for donors 1 and 2 but not for donor 3 on day 15.

Since the NK cell growth rate was mainly different between days 3 and 8 (Figure 1D), understanding the phenomena, for which subclusters CD8^+^ Tm-2 to CD8^+^ Tm-5 seemed to be important on these days, could elucidate potential causes for the growth difference. Figure 4G provides a comparison of the gene expression of these subclusters for the all donors and time points and the accompanying Table 3 categorizes the functions of a selection of the differentially expressed genes.

**Table 3:**
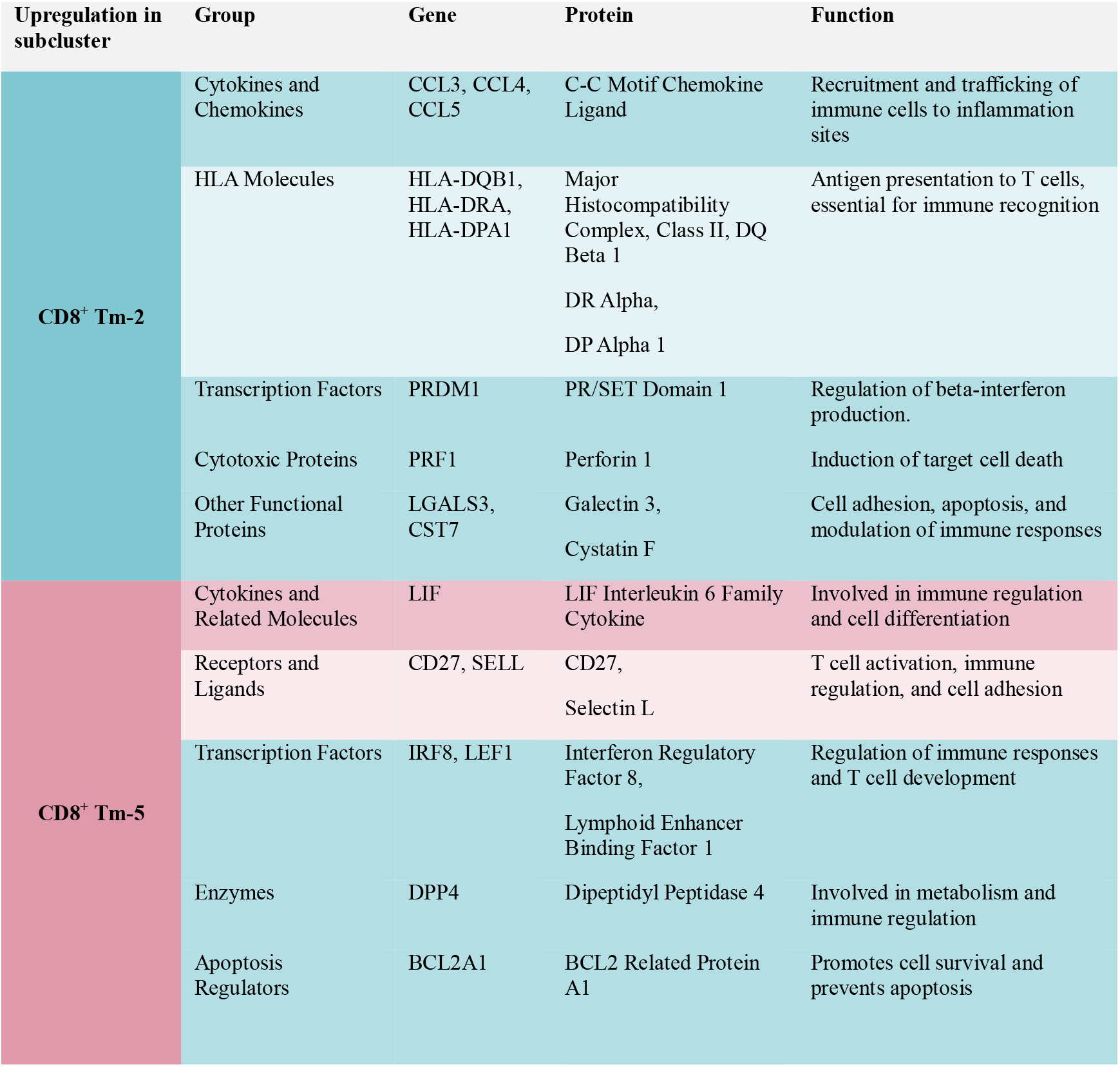
Gene descriptions selected significantly differentially expressed genes between subclusters CD8^+^ Tm-2 and CD8^+^ Tm-5 for all the donors and days.

Subcluster CD8^+^ Tm-2 displayed an increased expression of CC chemokines, specifically CCL3, CCL4, and CCL5. These typically act as chemo-attractants, by binding to the CCR5 receptor on immune cells including NK cells, with CCL5 known to specifically attract CD56^bright^ NK cells [46]. Subcluster CD8+ Tm-2 exhibited upregulation of cytotoxic proteins, such as *perforin 1* (PRF1), while subcluster CD8+ Tm-5 showed increased expression of CD27, indicative of a less differentiated state [47]. Finally, IRF8, a transcriptional regulator of interferon gamma (IFNγ) production was upregulated in the CD8^+^ Tm-5 subcluster. The effect of this upregulation was corroborated in the IFNG mRNA expression displayed in the violin plot in Figure 5C.

**Figure 5:**
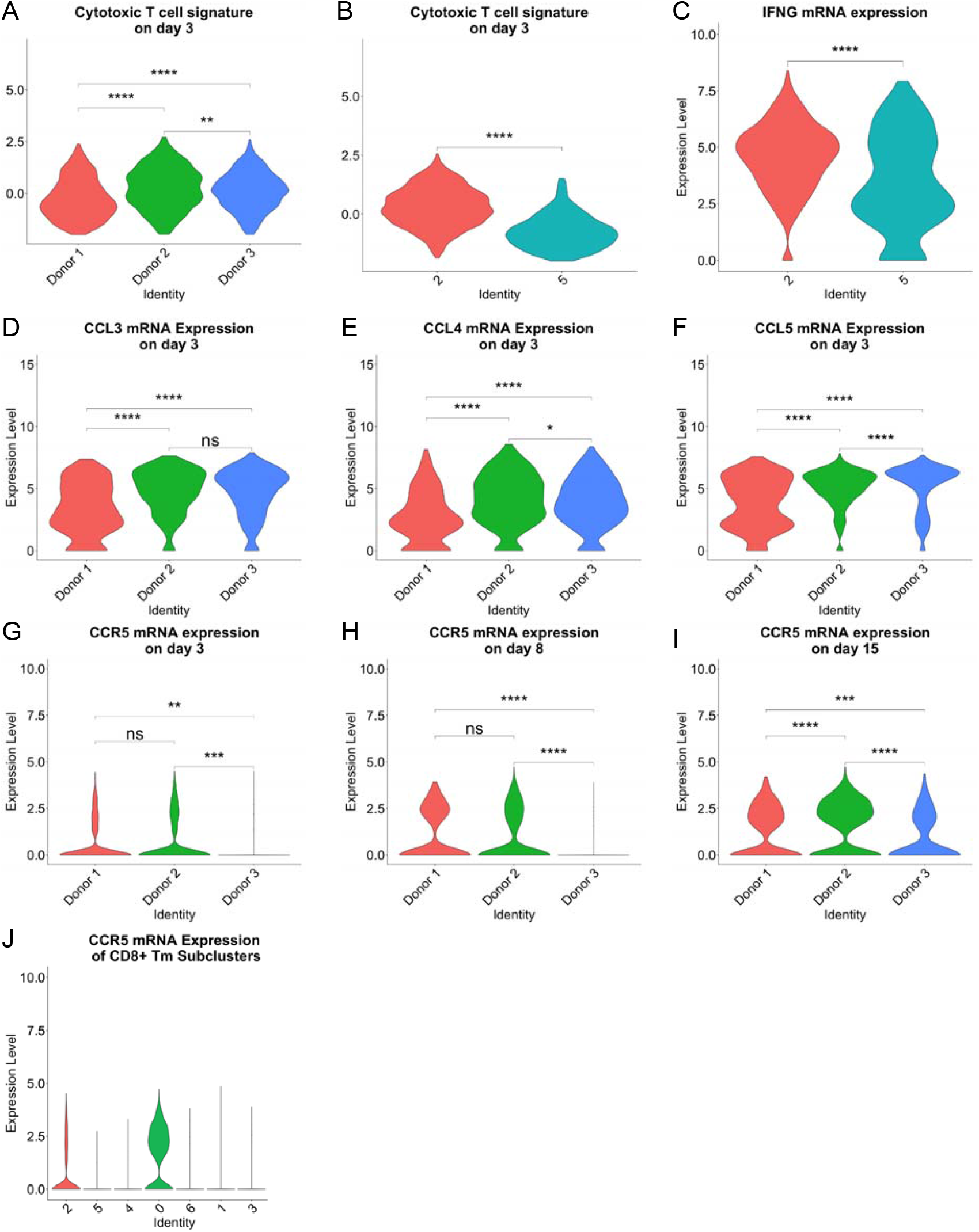
A) Cytotoxic T cell signature of the CD8^+^ Tm cells from the different donors on day 3. B) Cytotoxic T cell signature for the two most prevalent CD8^+^ Tm subclusters (CD8^+^ Tm-2 and CD8^+^ Tm-5) on day 3. C) Interferon gamma expression in the CD8^+^ Tm subclusters 2 and 5. D-F) mRNA expression of the cytokines CCL3, CCL4, CCL5 on day 3 for CD8^+^ Tm cells of the different donors. CD8^+^ Tm cell mRNA expression of the CCR5 chemokine receptor on G) day 3,H) day 8, I) day 15, and J) at all the days for each subcluster.

The significant upregulation of all these genes indicated that CD8^+^ Tm-2 subcluster was more cytotoxic than CD8^+^ Tm-5 subcluster. Comparing the cytotoxic T cell signature (NKG7, CCL4, CST7, PRF1, GZMA, GZMB, IFNG, CCL3) in Figure 5B between these two CD8^+^ Tm subclusters confirmed that CD8^+^ Tm-2 subcluster was indeed more cytotoxic than CD8^+^ Tm-5 subcluster. As previously observed, compared to donors 2 and 3, on day 3 the percentage of the CD8^+^ Tm-5 subcluster was highest in donor 1 while CD8^+^ Tm-2 subcluster was smaller (Figure 4 A-F). This was aligned with the results of Figure 5A, showing that the cytotoxic T cell signature for donor 1 was significantly lower than donors 2 and 3.

The temporal patterns of the CD8^+^ Tm subcluster development showed as well that the difference among the donors in the ending CD8^+^ Tm cell percentage was largely driven by the subcluster 0 evolution, Figures 4A – 4F. The donors that had a low ending NK cell percentage, donors 1 and 2, had a high CD8^+^ Tm cell percentage of more than 40%, while for donor 3 this percentage was around 10%, Figures 4A – 4C. The high ending percentage of NK cells of donor 3 could thus be driven by the lack of expansion of the CD8^+^ Tm cells, in particular subcluster 0.

Figures 5D – 5F show the mRNA expression levels of the chemokines CCL3, CCL4, and CCL5 on day 3. All these chemokines bind to the receptor CCR5, and activation of this receptor has been indicated to promote T cell growth [48]. Donors 2 and 3 had a high expression of these chemokines, which was expected due to the cytotoxic CD8^+^ Tm-2 phenotype representing a large fraction on day 3. However, the chemokine receptor CCR5 mRNA was not detected in CD8^+^ Tm cells from donor 3 culture on days 3 and 8, and significantly reduced on day 15 (Figure 5G – 5I). Finally, Figure 5J reveals that subcluster CD8^+^ Tm-0 was the primary contributor for the expression of CCR5 mRNA for all the donors analyzed together.

## Discussion

The consistent manufacturing performance of *ex vivo* expanded cells is of paramount importance for the future of cell-based therapies [49]. The aim of the current study was to investigate the dynamics of an *ex vivo* feeder cell free NK cell expansion used for an autologous cell therapy, and to identify key targets that provided a high likelihood of uncovering the donor-to-donor heterogeneity in this process. This was accomplished by a combination of flow cytometry and single cell transcriptomics. The flow cytometry was first used to identify key time points, i.e., points where the culture performances of the cultures differed with the donors. Then these time points were selected for analyses by single cell transcriptomics.

The three donors used in this study had a wide range of NK cell fold change and ending purity. Donors 2 and 3 had high fold expansion and donor 3 had high ending purity, making donor 3 the best performing donor. The fold change in NK cell numbers between donors differed most notably between days 3 and 8, with donor 1 exhibiting a slower growth rate compared to the other donors. This slow growth rate caused the lower fold expansion of donor 1 and therefore, days 3 and 8 represented the most interesting samples to analyze by scRNA-seq, supplementary to the starting and end time points, days 0 and 15 respectively.

After the scRNA-seq data had been aligned and filtered, the lack of clustering by donor shows limited batch effects. The cell type annotation, performed by the pipeline provided by BD™, showed a good agreement to the cell types determined by flow cytometry, except however for day 0, where the scRNA-seq predicted a higher proportion of NK cells than the flow cytometry. It has been shown previously that scRNA-seq predicts a lower proportion of NK and T cells and a much higher proportion of monocytes compared to flow cytometry [50]. The particular flow cytometry panel used in this study first gated out any monocytes and B cells. On day 0 there was a large fraction of monocytes detected by scRNA-seq, >50%, so upon removal of this cell type the proportions of NK and T cells increased dramatically, surpassing the results from flow cytometry. After day 0, when the monocyte population was very low, the flow cytometry and scRNA-seq cell type predictions agreed very well with each other.

To decipher the information about phenotypes within each individual cell type, subclustering was performed. The clustering algorithm was run multiple times, and, after each run, the silhouette score, which provides a performance metric for the clustering, was calculated. Based on this score the optimal parameters for the subclustering were chosen. This allowed for a data-driven approach of the parameter selection, which eliminated the normally subjective nature of clustering. The results of this subclustering revealed many distinct groups within each cell type. These groups, i.e., subclusters, were then annotated using 6 additional databases, beyond the one used in the primary cell type annotation. The additional databases largely agreed in the predicted cell types, except for some subclusters and cell types such as dendric cells, which mostly annotated to monocytes. However, given that these cell types share the same progenitor cell, myeloid cells, this overlap was perhaps not unexpected. A subcluster of CD8^+^ Tm cells, CD8^+^ Tm-0, was annotated to NK-like T cells in the only database that contained a signature for NK-like T cells. This subcluster had markedly different behavior in donor 3 culture, which ended with a high NK cell fraction, suggesting that NK-like T cells might play a pivotal role in the ending purity of *in vitro* NK cell expansion.

Compared to each other’s, the NK cell subclusters of the high-expansion donors, 2 and 3, showed a larger similarity over time than the low-expansion donor 1, with a key difference being, for this latter, the occurrence of subcluster NK-5 observed after day 3 and increase until day 15 (Figure 2D). Despite a significant fraction of the total NK cells being NK-5 subcluster cells in donor 1, the NK cell apparent growth rates, measured by flow cytometry, showed only a small difference between the donors from day 8 to day 15. This suggests that subcluster NK-5 did not detrimentally affect the growth of the NK cells in donor 1, but rather that the occurrence of subcluster NK-5 was an effect of an event that happened between days 3 and 8. The trajectory inference of the NK subclusters, Figure 2G, shows that subcluster NK-5 resulted from a branching between subclusters NK-3 and NK-0. This further indicates that the event that caused the occurrence of subcluster NK-5 happened at the same time as the cells were differentiating into cluster NK-3, i.e., between days 3 and 8. The NK cells of donor 1 were found to have upregulation in certain inflammatory and stress related genes (i.e., CASP5, IL18RAP, FOSB, RORA) on day 3 compared to the other donors, Figure 3A and Table 1. This suggests that the heterogeneity among the donors was caused by an event which caused a more inflammatory and stressful environment in the culture of donor 1 compared to the other donors. A recapitulation table is given in Table 4 listing the major observations of the study.

**Table 4:** Recapitulation of the major comparative observations of the study, where lower values are highlighted in blue and higher in pink.

| Factors | Donor 1 | Donor 2 | Donor 3 | Figures & Tables |
| --- | --- | --- | --- | --- |
| Ending NK cell purity | 13.7% | 26.1% | 80.5% | Table 1 |
| Fold expansion of NK cell | Low | High | High | Table 1 |
| Upregulation in certain inflammatory and stress related genes (i.e., CASP5, IL18RAP, FOSB, RORA) on day 3 | Higher | Lower | Lower | Figures 3A & Table 1 |
| Inflammatory and stressful environment | More | Less | Less |  |
| Subcluster NK-5 observed after day 3, with occurrence due to event happening between day 3 and 8 | Increasing | Flat & low | Flat & low | Figure 2 |
| Scores for the CD56<sup>Bright</sup> CD16<sup>-</sup> phenotype on day 3 and day 8 | Low | High | High | Figures 3B – 3G |
| Scores for the CD56<sup>Dim</sup> CD16<sup>+</sup> phenotype on day 3 and day 8 | High | Low | Low | Figures 3B – 3G |
| Expansion of the CD8<sup>+</sup> Tm cells | High | High | Low | Figures 4A-C |
| CD8<sup>+</sup> Tm-2 subcluster on day 3 | Less prevalent | More prevalent | More prevalent | Figures 4A-F |
| Chemokine and other cytotoxic genes of CD8<sup>+</sup> Tm-2 subcluster |  | Upregulation | Upregulation | Figures 5B, 5D-F |
| CD8<sup>+</sup> Tm-5 subcluster on day 3 | More prevalent | Less prevalent | Less prevalent | Figures 4A-F |
| Genes of CD8<sup>+</sup> Tm-5 associated with a less differentiated and less cytotoxic phenotype | Upregulation |  |  | Figure 5 & Table 3 |
| Expansion of the CD8<sup>+</sup> Tm-0 subcluster annotated as NK-like T cell | High | High | Low | Figures 4D – 4F & Figure S1 |
| CCR5 mRNA by CD8<sup>+</sup> Tm cells on day 3 & 8 (activation of CCR5 known to aid T cell growth) | Observed | Observed | Absent | Figures 5G – 5J |
| CCL3, CCL4, and CCL5 mRNA on day 3 | Low | High | High | Figures 5D-F |

Compared to donor 1, the lower levels of inflammatory and stress signaling observed in the NK cells of donors 2 and 3 likely favored a shift towards a more proliferative phenotype. This can be seen when comparing the gene signatures, in Figures 3B – 3G, for the two main phenotypes of NK cells, CD56^Bright^ CD16^−^ and CD56^Dim^ CD16^+^. The cells issued from donors 2 and 3 had higher scores for the CD56^Bright^ CD16-phenotype on day 3 and even more on day 8, as well as lower scores for the CD56^Dim^ CD16^+^ phenotype on the same days. In addition, donor’s 2 and 3 cells had also a lower expression of CD16 mRNA than the ones of donor 1. It has been shown that the CD56^Bright^ CD16^−^ NK cell phenotype has a greater proliferative capacity than its CD56^Dim^ CD16^+^ counterpart [44]. Possibly the stress and inflammatory signals present between day 3 and day 8 for donor 1 caused a slower shift from the CD56^Dim^ CD16^+^ to the CD56^Bright^ CD16^−^ phenotype. This delayed transition could also have impacted the proliferation rates during this same time frame.

As observed in Figure 2D – 2F, the major ending subcluster of NK cells was NK-0. It has been previously reported that the *ex vivo* expansion protocol used in this work leads to a unique phenotype of NK cells [45]. The genes encoding the surface proteins making up this phenotype, Table S1, were used to create an *ex vivo* NK cell signature, confirming the predominance of NK-0 subcluster, Figure 2H. This signature correlated well with the pseudotime showing a greater intensity the farther a cell progressed along its trajectory. While NK-0 represented the largest subcluster of NK cells at day 15 several other subclusters were also present. Further investigation into the clinical effectiveness of each of these NK subclusters would be interesting to study to evaluate the potential to improve the efficacy of the treatment by modifying the production process to increase the fraction of a particular subcluster.

The heterogeneity among the NK cells was larger between day 3 and day 8, as evidenced by the appearance of the NK-5 subcluster and the upregulation of the inflammatory and stress response genes in donor 1. This however, only indicated an effect and not the root cause, for which the CD8^+^ Tm cells could provide a clue. Similarly, to the NK cells, the subcluster fractions of CD8^+^ Tm cells changed over time, Figures 4A – 4F. Interestingly, on day 3, there was a substantially different distribution of the CD8^+^ Tm subclusters for the low-expansion donor 1 NK cells compared to the other donors (Figure 4D), with CD8^+^ Tm-5 subcluster being much more prevalent than the other donors. Figure 4G shows the differential gene expression of the two most prevalent CD8^+^ Tm subclusters on day 3, CD8^+^ Tm-2 and CD8^+^ Tm-5. The CD8^+^ Tm-2 subclusters showed upregulation of chemokine and other cytotoxic genes, whereas CD8^+^ Tm-5 showed an upregulation in genes associated with a less differentiated and less cytotoxic phenotype. The CD8^+^ Tm-2 subcluster had a significantly higher score than the CD8^+^ Tm-5 subcluster in the gene signature for cytotoxic T cells. This could indicate that the additions of anti-CD3 (OKT-3) and/or IL2 were less effective or insufficient in donor 1 culture compared to the other donors. The addition of these factors in the culture induces the secretion of cytokines by the T cells, which is known to help the NK cells growth and activation. This suggests that these factors might be needed in different amount for different donors, appealing for a donor specific tuning of their supplementation. It could also be interesting to monitor the activation status of the CD8^+^ Tm cells early in the culture after anti-CD3 stimulation, and if necessary (as it could have been the case for donor 1), re-administer OKT-3.

The chemokines CCL3, CCL4, and CCL5, which are known to be important for the recruitment of NK cells [46], were upregulated in donors 2 and 3 cell cultures on day 3. In particularly, CCL5 has been shown to attract CD56^Bright^ NK cells. It can be hypothesized that this upregulation caused an environment more favorable, i.e., with a lower stress level, for a shift towards the CD56^Bright^ phenotype observed in donors 2 and 3. In the future, one could explore if an addition of these chemokines could ensure a high fold expansion of the NK cells in all the cultures regardless of the T cell activation status.

The observation of the temporal pattern of the CD8^+^ TM subclusters gave also a clue of the ending purity of NK cells for each donor. Donors 1 and 2 (13.7% and 26.1%) had low ending NK cell purity and donor 3 a high purity (80.5%), see Table 1. A primary difference between these two groups was the expansion of the CD8^+^ Tm-0 subcluster, low for donor 3 cells and high for the other donors, see Figures 4D – 4F. As mentioned above, this subcluster was also annotated as an NK-like T cell by the only database that contained the signature for this cell type, see Figure S1. Another unique feature, for which these two groups diverged, was the expression of CCR5 mRNA by CD8^+^ Tm cells, observed for donors 1 and 2 but absent or weak for donor 3, see Figures 5G – 5J. It is known that the CCR5 activation is implicated in aiding T cell growth [48]. Importantly, this difference was observed already at day 3 before the CD8^+^ Tm-0 subcluster started a major expansion for donors 1 and 2. This indicates that the lack of CCR5 expression for the CD8^+^ Tm cells of donor 3 was associated with the very low expansion of this subcluster in comparison with donors 1 and 2. This implies as well that CCR5 could be used as a marker to gauge the proliferative capacity of the CD8^+^ Tm cells during a cultivation process, potentially giving the operators an early warning of a low NK cell purity batch. It could also suggest an interesting signaling pathway to target, for instance by the addition of blocking peptides in order to halt the growth of CD8^+^ Tm cells, thereby increasing the ending purity of NK cells.

## Conclusion

This study reveals the dynamics of an *in vitro* culture of PBMCs with the aim of providing a high purity and expansion of NK cells for an autologous cell therapy product. Through the analyses of unique phenotypes and subclusters within both the NK and CD8^+^ Tm cells, as well as their correlation to the cell expansion process, we provided key insights into the donor-related variation of this process outcome. While the limited number of donors used here somewhat limited the statistical power of this study, the insights into the dynamics of the culture process identified several phenomena, which will be important to explore in future studies. Further investigations into the roles that the CCL chemokines play in the expansion process of NK cells in an *in vitro* system could yield significant improvements to this process, as well as, investigating the role of CCR5 on the expansion of CD8^+^ Tm cells to improve the purity of the final product. Tuning of the protocol for the supplementations of the anti-CD3 antibody, OKT3, and/or IL2 in a donor specific way could also potentially improve this outcome, and could be explored based on donor-specific markers.

Importantly, these findings highlight the role that single cell transcriptomics can play in the process development for a cell therapy product by providing a limited list of potentially influential components, important biomarkers and/or process changes. This approach can greatly enhance the development time, the process performance and the cell product quality, generating *in fine* an improved product for the patients.

## Declaration of competing interest

None of the co-authors have any conflicts of interest to declare for the project.

## Funding

This research was supported by the Competence Centre for Advanced BioProduction by Continuous Processing, AdBIOPRO, funded by Sweden’s Innovation Agency VINNOVA (diaries nr. 2022-03170), XNK Therapeutics (XNK Therapeutics has since ceased operations) and partially supported by Digital Futures, KTH Royal Institute of Technology.

## Author Contributions

Conceptualization, B.L., P.H., P.B., V.C.

Methodology, B.L., P.H., M.Z., S.R.

Software, B.L., M.Z.

Data curation, B.L., M.Z.

Investigation, B.L., P.H.

Validation, B.L., S.R., P.H.

Formal analysis, B.L.

Supervision, V.C., P.H.

Funding acquisition, V.C., P.B.

Visualization, B.L.

Project administration, V.C., P.H.

Resources, V.C., P.B

Writing—original draft preparation, B.L.

Writing—review and editing, B.L., P.H., S.R., M.Z. V.C.

All authors have read and agreed to the published version of the manuscript.

## Notes

### Competing Interest Statement

The authors have declared no competing interest.

