## Supplement for "Single-Cell Transcriptomics Reveals Dynamics of NK Cell Expansion in a Feeder Cell-Free Culture of PBMCs - Implications for Immunotherapy"

a

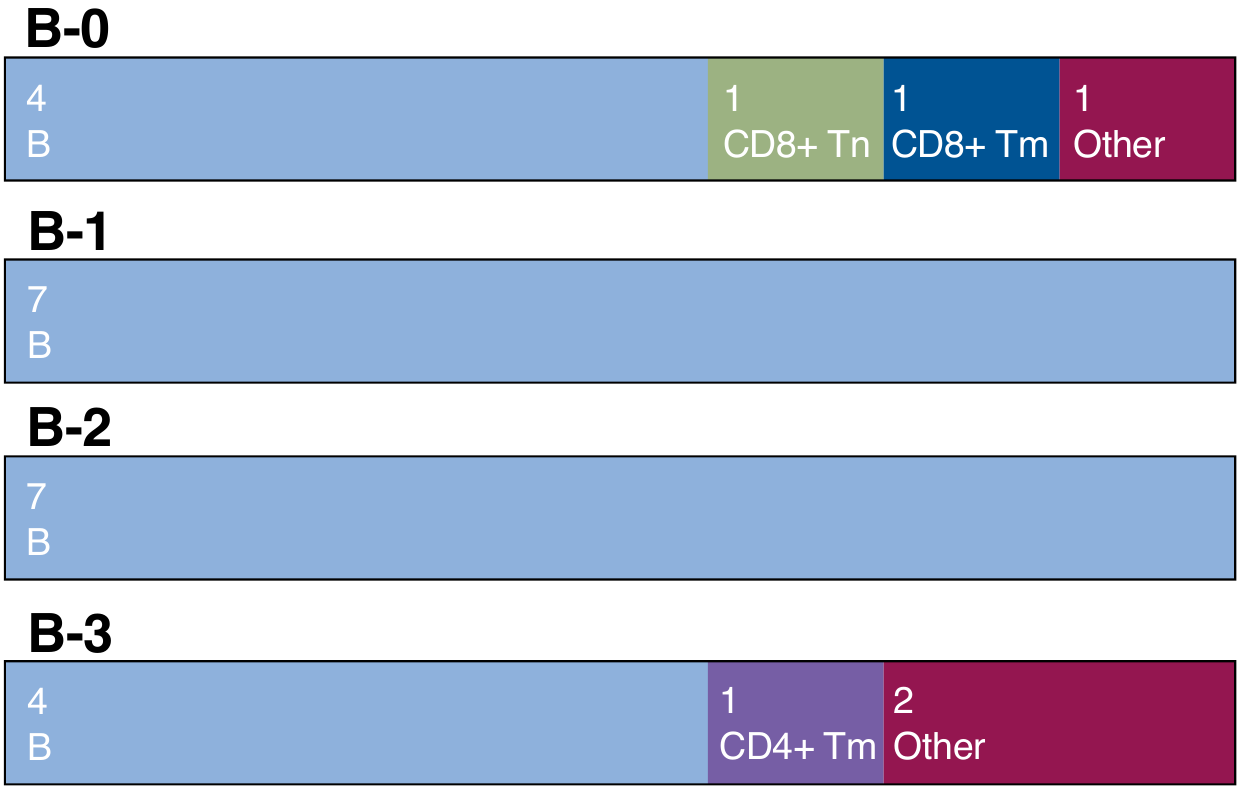

b

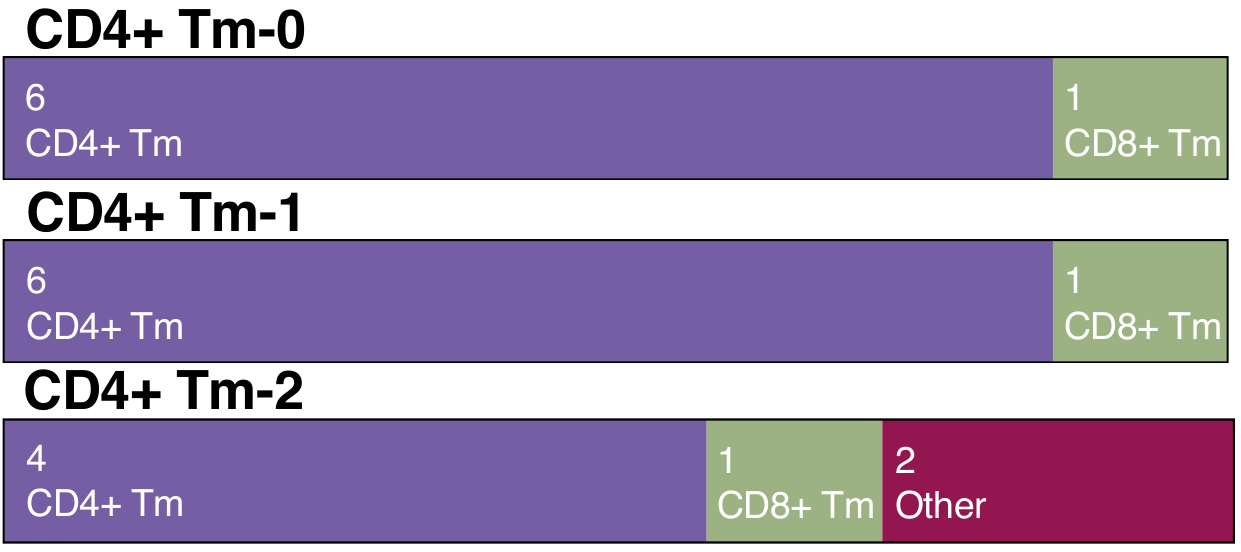

c

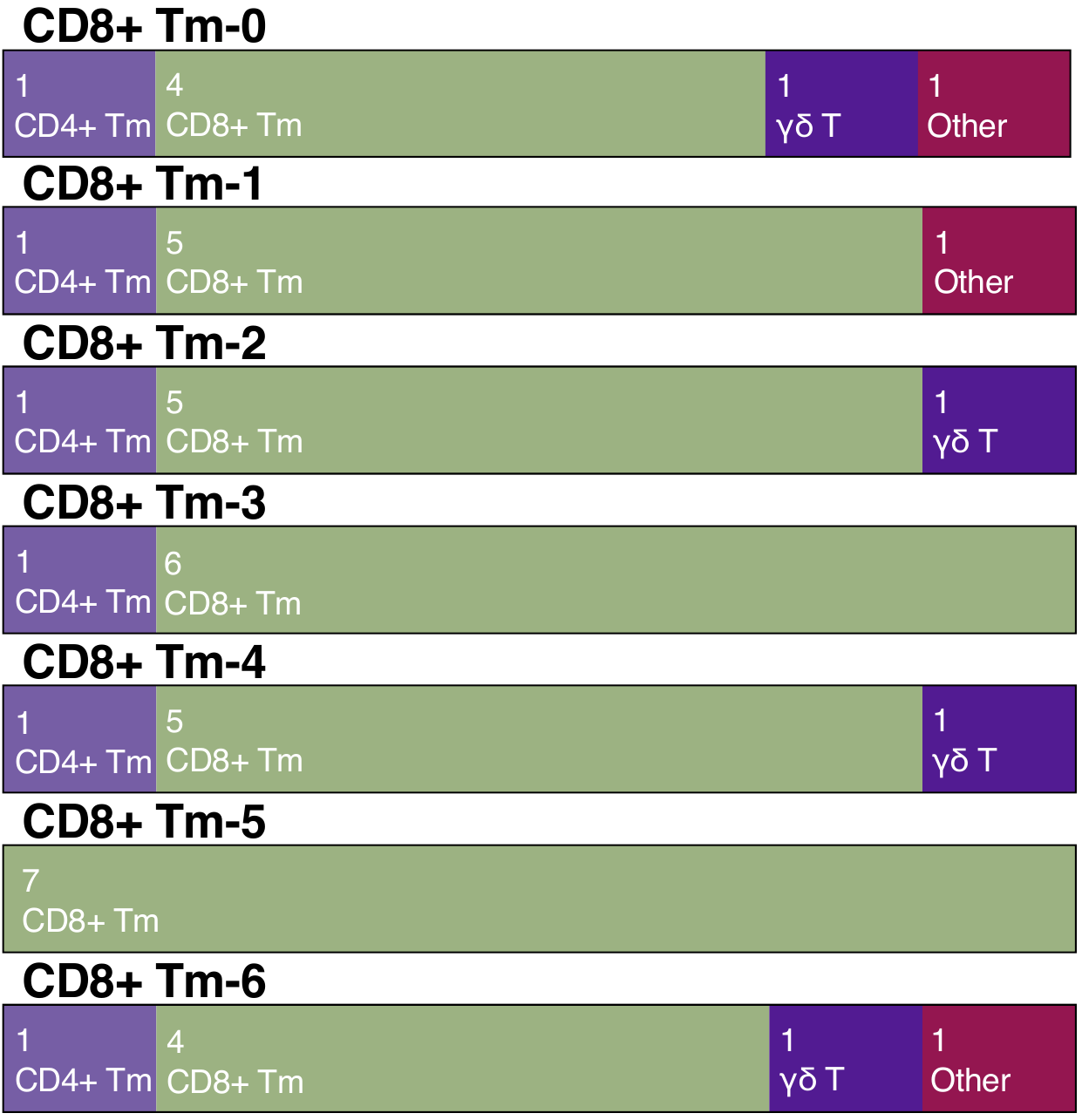

d

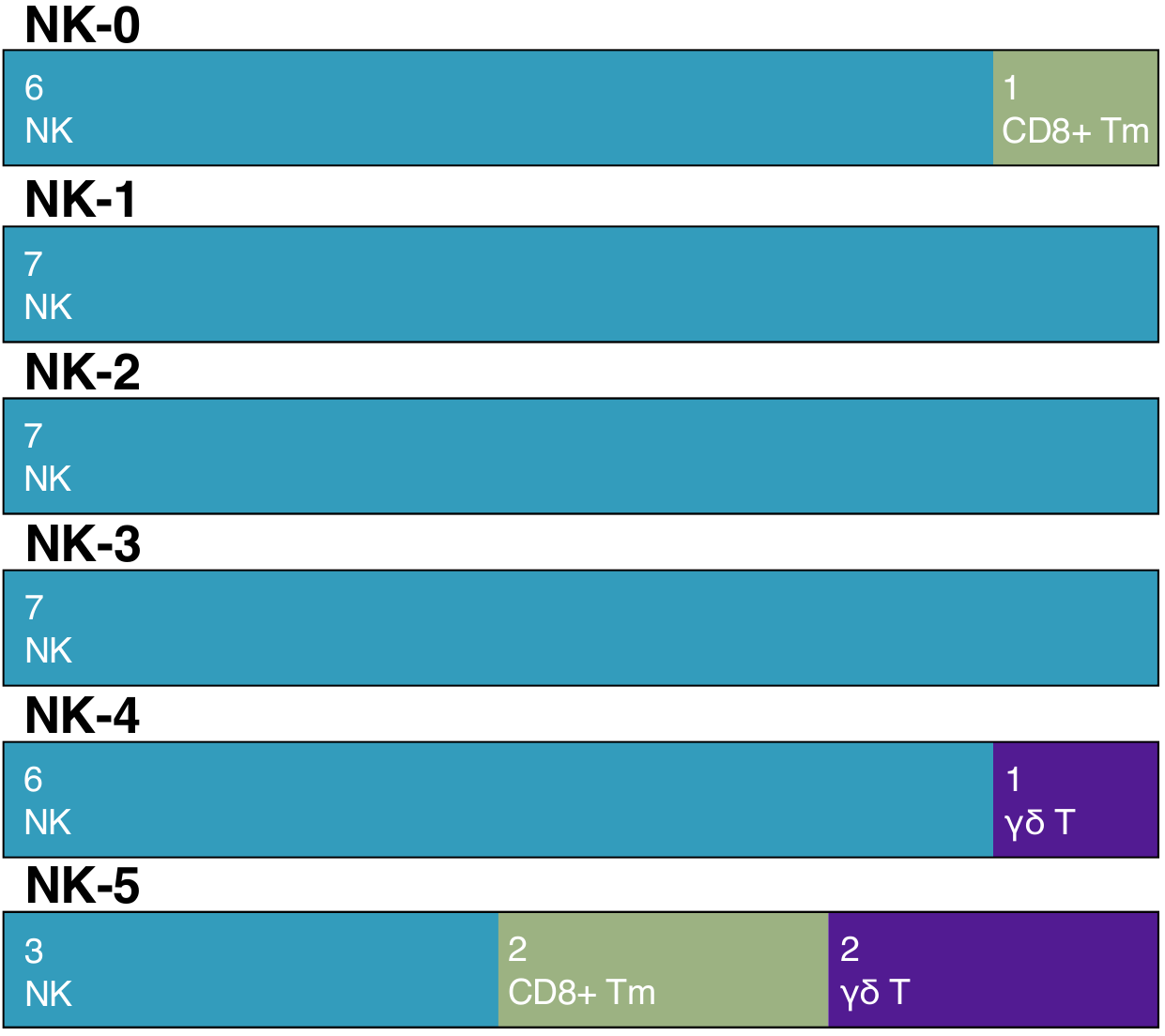

**Figure S1. Consensus cell type prediction across seven databases.**
This figure illustrates the agreement among the seven databases (MID,HPCA,sc-Type,BE, DICE, azimuth, and the BD Sequence Analysis Pipeline) on the predicted cell type identities for a) B cells, b) CD4+ Tm cells, c) CD8+ Tm cells, d) NK cells. Highlighting shared classifications and areas of divergence. The consensus underscores the reliability of the predictions and provides insight into the robustness of cell type annotations in the dataset.

a

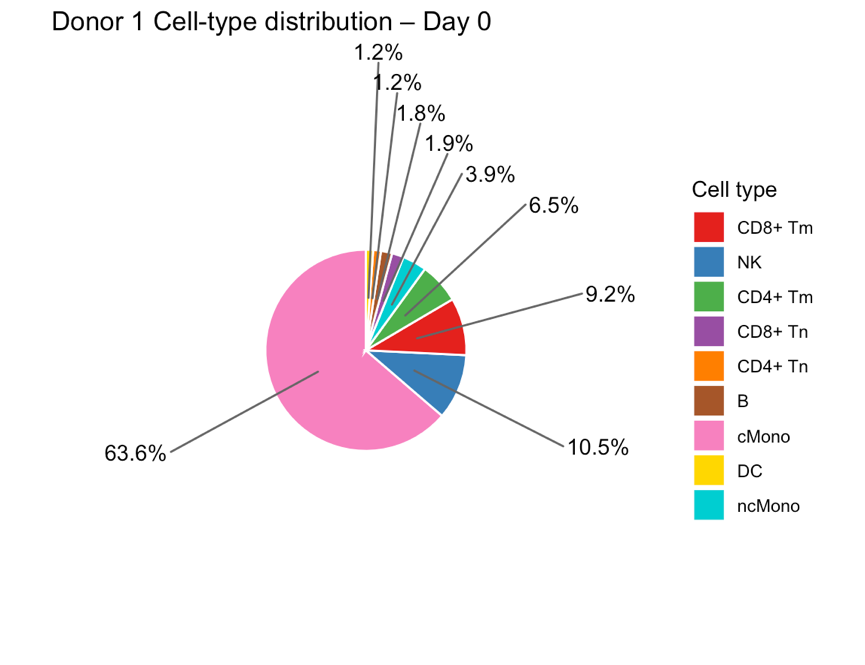

b

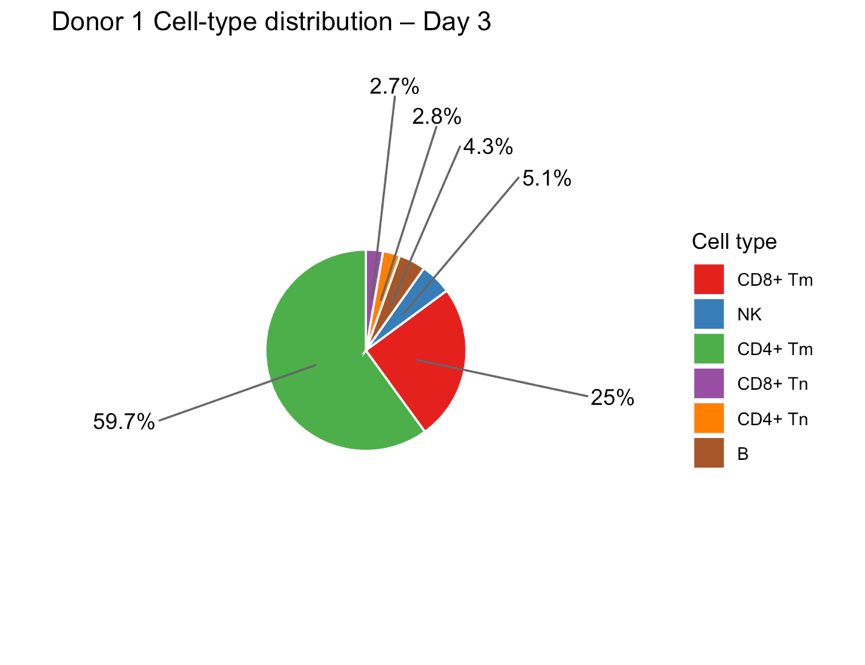

c

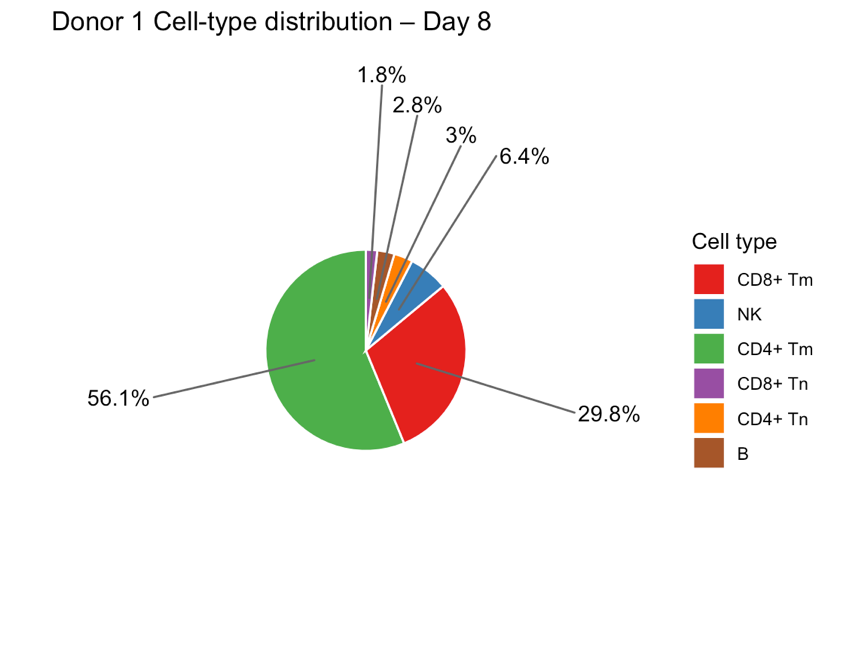

d

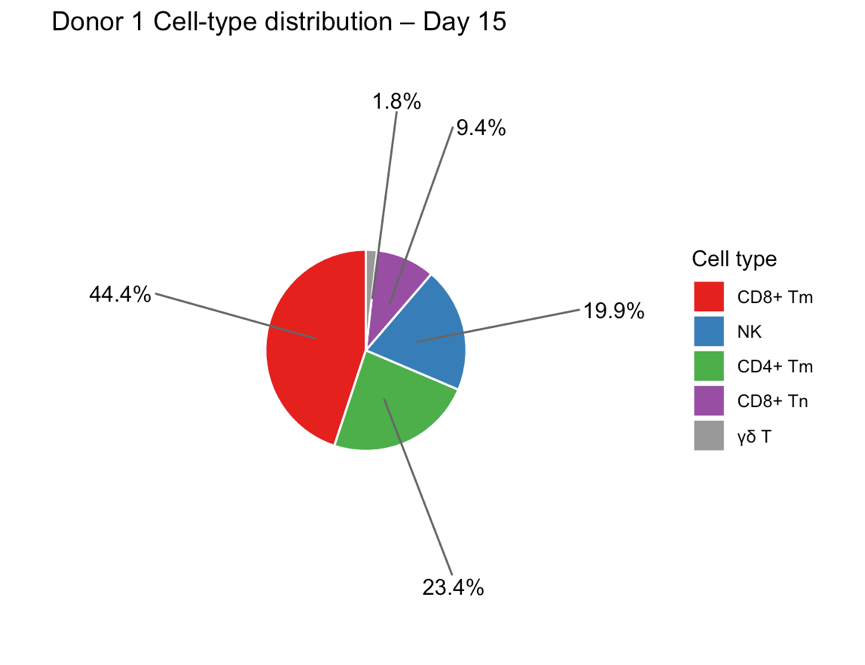

**Figure S2. Cell type composition during ex vivo expansion over time for donor 1.**

Pie charts showing the percentage distribution of major cell types at four time points during the expansion protocol: Day 0 (baseline), Day 3, Day 8, and Day 15. Each chart displays the relative proportions of each cell type as percentages of the total cell population. Cell types representing <1% of the total population are excluded from visualization for clarity.

a

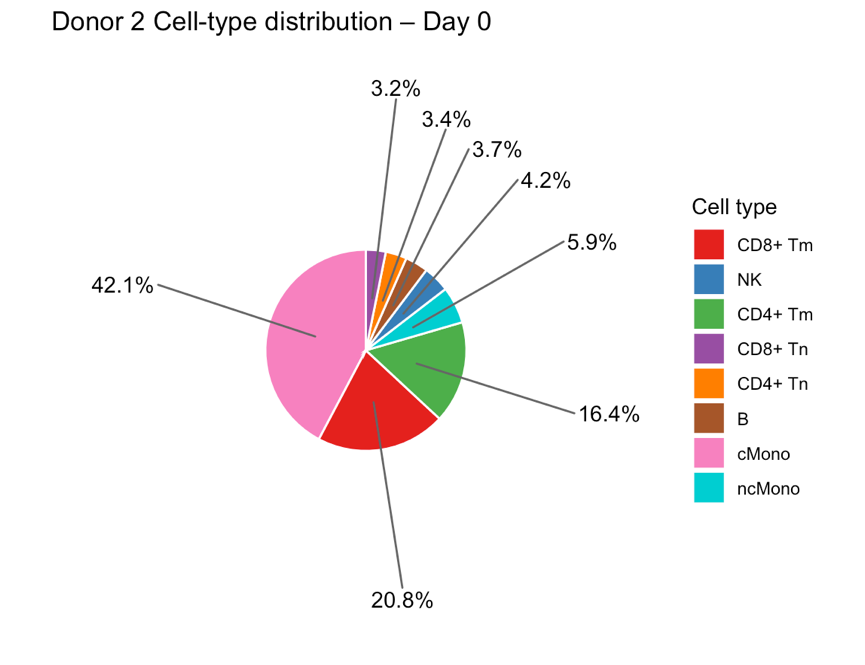

b

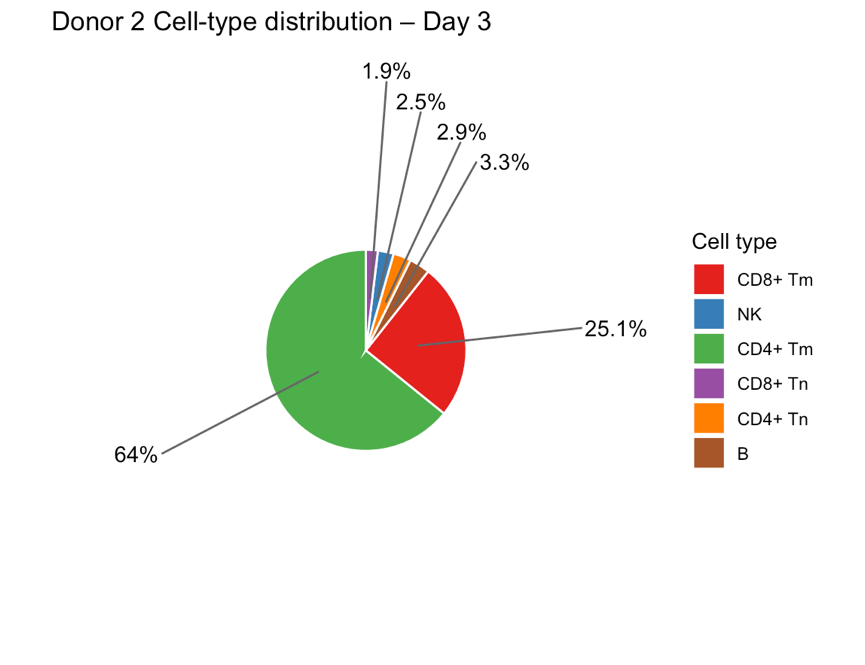

c

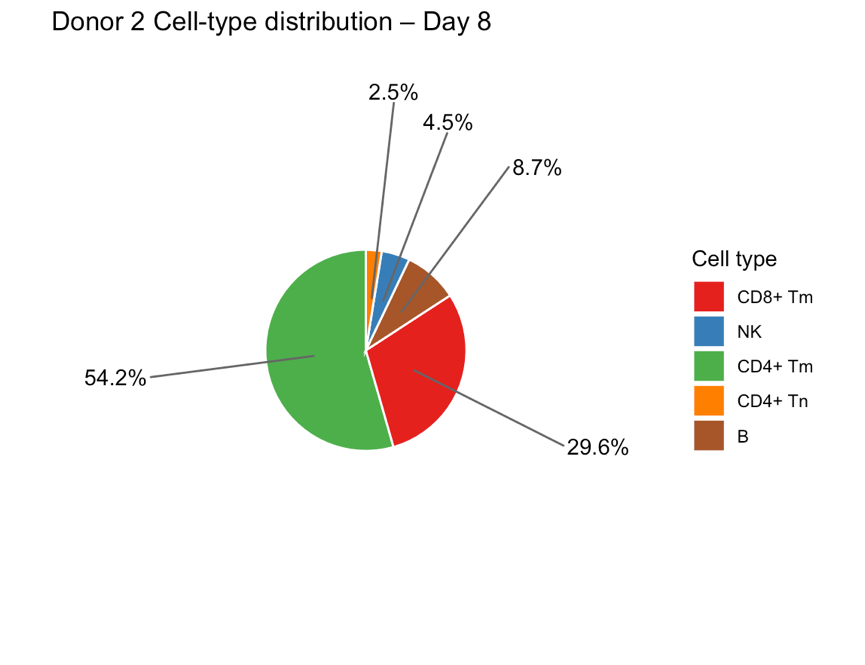

d

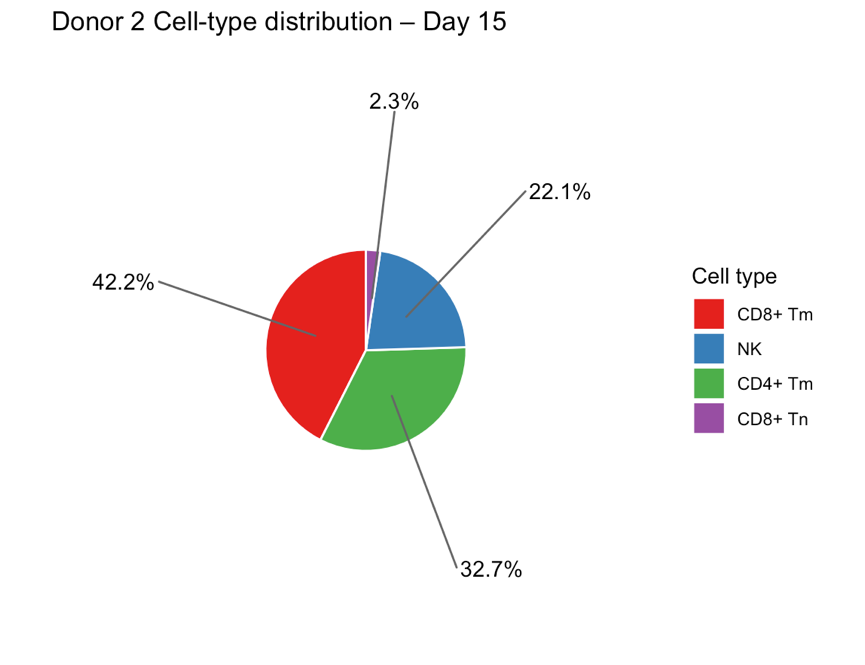

**Figure S3. Cell type composition during ex vivo expansion over time for donor 2.**

Pie charts showing the percentage distribution of major cell types at four time points during the expansion protocol: Day 0 (baseline), Day 3, Day 8, and Day 15. Each chart displays the relative proportions of each cell type as percentages of the total cell population. Cell types representing <1% of the total population are excluded from visualization for clarity.

a

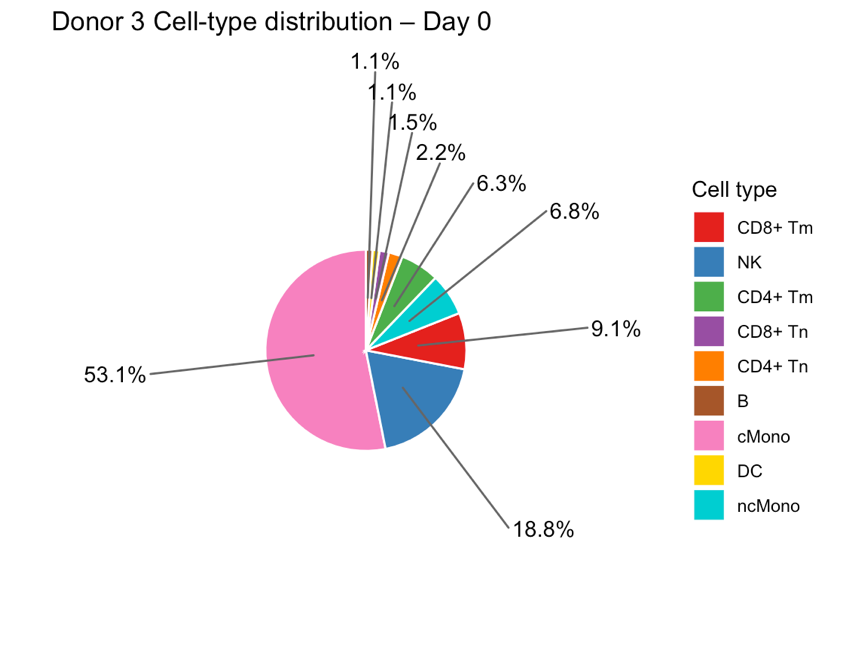

b

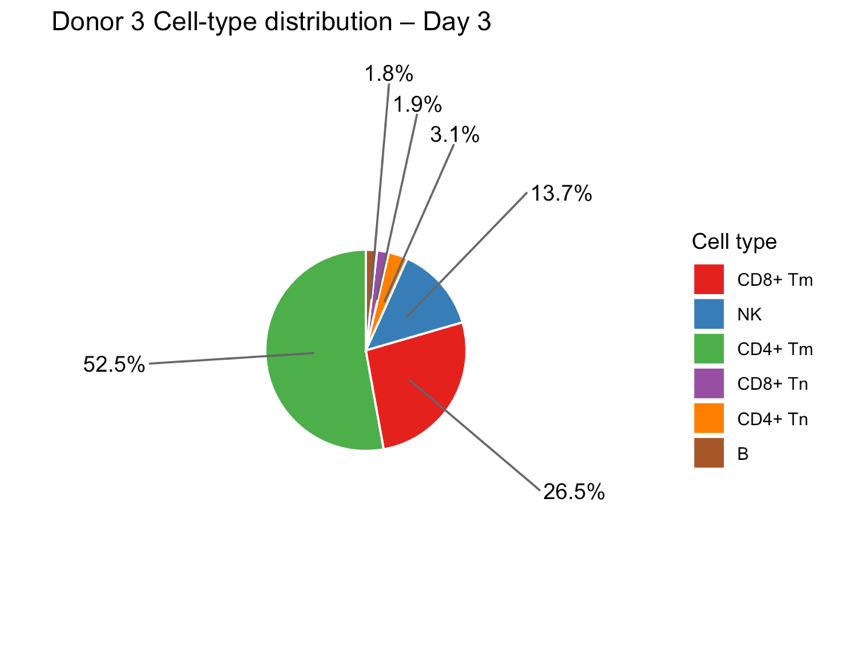

c

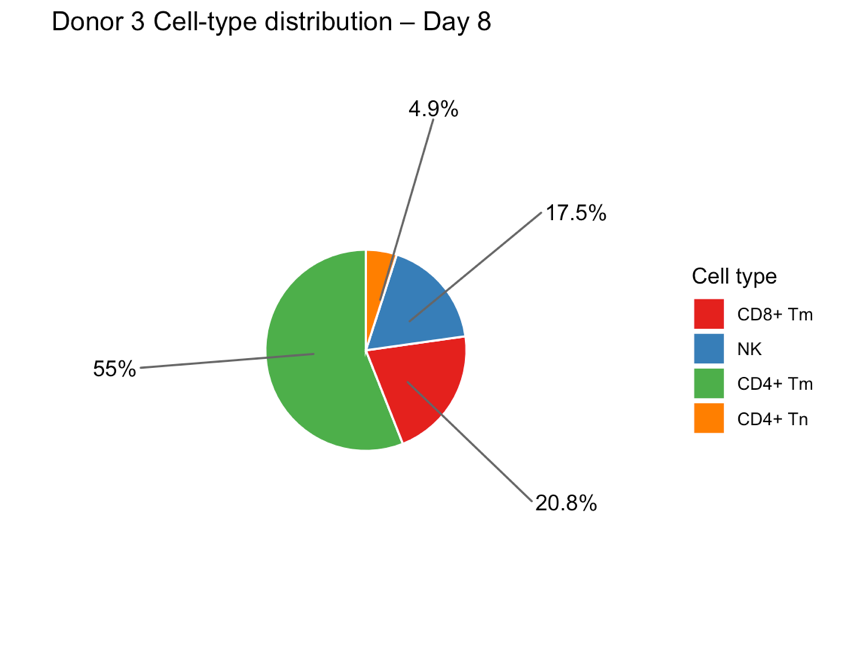

d

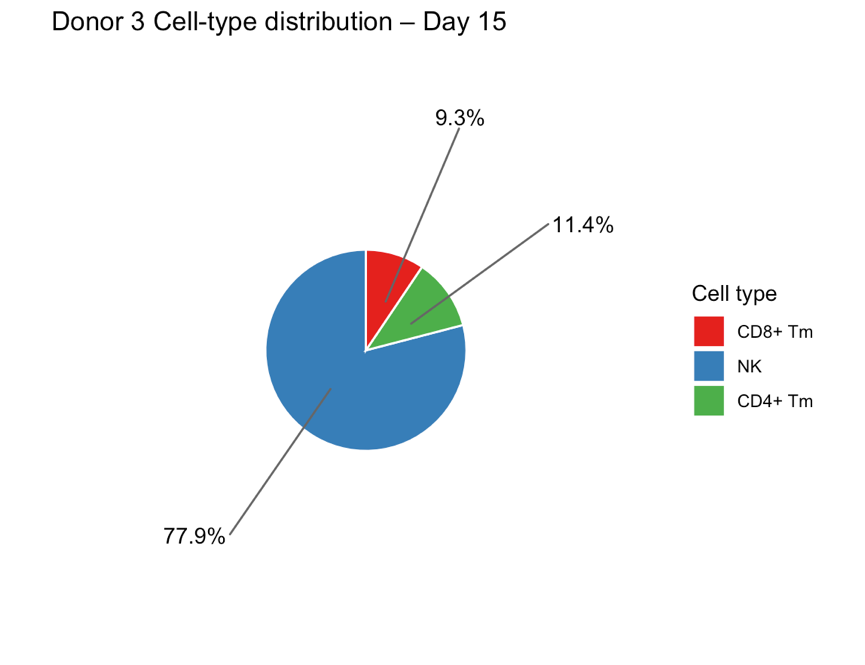

**Figure S4. Cell type composition during ex vivo expansion over time for donor 3.**

Pie charts showing the percentage distribution of major cell types at four time points during the expansion protocol: Day 0 (baseline), Day 3, Day 8, and Day 15. Each chart displays the relative proportions of each cell type as percentages of the total cell population. Cell types representing <1% of the total population are excluded from visualization for clarity.

| CD56 Bright | CD56 Dim | Ex vivo Phenotype |
| --- | --- | --- |
| IL23R | CMKLR1 | NCAM1 |
| RORC | CX3CR1 | MKI67 |
| DPP4 | LAIR2 | FCGR3A |
| CXCR6 | FCGR3A | HLA.DRA |
| KIT | CXCR1 | KLRK1 |
| CCR7 | KIR2DL1 | CD244 |
| CCR8 | CXCR2 | HAVCR2 |
| LAG3 | GZMB | TIGIT |
| FLT3 | B3GAT1 |  |
| IL7R | PI3 |  |
| CCR5 | GZMH |  |
| GZMK | HAVCR2 |  |
| CD33 | PRF1 |  |
| PDCD1 | NKG7 |  |
| CCR2 | TBX21 |  |
| CD5 | NCR3 |  |
| LIF | ARL4C |  |
| JCHAIN | CD247 |  |
| CD8B | CST7 |  |
| TRAT1 | SELPLG |  |
| CD3D | CHI3L2 |  |
| CD28 | ITGAM |  |
| CD27 | GZMA |  |
| RNASE2 | CD244 |  |
| NRP1 | ITGB2 |  |
| POU2AF1 | IFNG |  |
| DUSP4 | KLRG1 |  |
| BLK | IL12RB1 |  |
| CCL20 |  |  |
| CD3G |  |  |
| YBX3 |  |  |
| MYC |  |  |
| LEF1 |  |  |
| IL2RA |  |  |
| IL1R2 |  |  |
| TNFSF8 |  |  |
| TLR2 |  |  |
| SNCA |  |  |
| TCF7 |  |  |
| CXCL1 |  |  |
| MZB1 |  |  |
| IL18 |  |  |
| CD3E |  |  |
| BIRC3 |  |  |
| IL1RL1 |  |  |
| ADGRG3 |  |  |
| CXCR3 |  |  |
| CXCL2 |  |  |
| ANXA5 |  |  |
| TNFSF13B |  |  |
| TCF4 |  |  |
| ENTPD1 |  |  |
| RGS1 |  |  |
| KLRC4 |  |  |
| LTB |  |  |
| CCR1 |  |  |
| S100A9 |  |  |
| MCM4 |  |  |
| KLRC1 |  |  |
| SELL |  |  |
| THBS1 |  |  |
| KLRC3 |  |  |
| CD44 |  |  |
| CD52 |  |  |
| CD2 |  |  |
| Table S1. Gene list for CD56 Bright and CD56 Dim NK cells from {Dogra, 2020 #207}. Gene list for Ex vivo Phenotype from {Nahi, 2022 #82}. | | |

| Difference in mean | NK-0 | NK-1 | NK-2 | NK-3 | NK-4 | NK-5 |
| --- | --- | --- | --- | --- | --- | --- |
| NK-0 | - | -0.96385 | -0.9464 | -0.45571 | -0.5811 | -0.55319 |
| NK-1 | 0.96385 | - | 0.01745 | 0.50813 | 0.38275 | 0.41065 |
| NK-2 | 0.9464 | -0.01745 | - | 0.49069 | 0.3653 | 0.3932 |
| NK-3 | 0.45571 | -0.50813 | -0.49069 | - | -0.12539 | -0.09748 |
| NK-4 | 0.5811 | -0.38275 | -0.3653 | 0.12539 | - | 0.02791 |
| NK-5 | 0.55319 | -0.41065 | -0.3932 | 0.09748 | -0.02791 | - |
| Table S2. Differences in module scores between NK cell subclusters for the *ex vivo* phenotype. This table shows the differences in module scores (reflecting the *ex vivo* phenotype of NK cells) between the subclusters. Each cell represents the difference between the module scores of the two subclusters indicated by the row and column. | | | | | | |

| Adjusted p-value | NK-0 | NK-1 | NK-2 | NK-3 | NK-4 | NK-5 |
| --- | --- | --- | --- | --- | --- | --- |
| NK-0 | - | 0 | 0 | 0 | 0 | 0 |
| NK-1 | 0 | - | 0.9734357 | 0 | 0 | 0 |
| NK-2 | 0 | 0.9734357 | - | 0 | 0 | 0 |
| NK-3 | 0 | 0 | 0 | - | 0.0010344 | 0.084686 |
| NK-4 | 0 | 0 | 0 | 0.0010344 | - | 0.9817327 |
| NK-5 | 0 | 0 | 0 | 0.084686 | 0.9817327 | - |
| Table S3. Adjusted p-values for comparisons of module scores between NK cell subclusters for the *ex vivo* phenotype. This table shows the adjusted p-values (from the function TukeyHSD using default parameters) for the comparisons of module scores between NK cell subclusters, assessing differences in the *ex vivo* phenotype of NK cells. The values indicate the statistical significance of the differences in module scores between the respective subclusters. | | | | | | |
